# An Innervated and Vascularized HNSCC-on-a-Chip Model Built on Defined and Tunable Engineered Extracellular Matrices

**DOI:** 10.64898/2026.01.16.699893

**Authors:** Alessia Maria Pignataro, Cornelia Christina Schwarz, Emilia Wiechec, Alessandro Cordiale, Shyama Sasikumar, Alexander Jenssen, Taka Ariyaberg, Lalit Pramod Khare, Ehsanul Hoque Apu, Karin Roberg, Sajjad Naeimipour, Gabriela Basile Carballo, Marcin Szczot, Daniel Aili, Marco Rasponi, Pierfrancesco Pagella

**Author notes:** corresponding author. **Corresponding author:** Pierfrancesco Pagella. these authors contributed equally to this work (Alessia Maria Pignataro, Cornelia Christina Schwarz).

## Abstract

Understanding the tumor microenvironment (TME) requires experimental platforms that faithfully recapitulate its key components. Here, we present an innervated and vascularized head and neck squamous cell carcinoma (HNSCC)-on-a-chip platform built with fully defined and tunable engineered extracellular matrices (eECMs). In a stepwise increase of complexity, we first co-cultured patient-derived HNSCC cells, cancer-associated fibroblasts, and endothelial cells within tailored eECMs, revealing matrix-dependent differences in self-organization and chemotherapeutic sensitivity. We then integrated these 3D constructs into a cancer–vasculature-interface, which enabled analysis of eECM-dependent directional collective migration and metastatization. Finally, we incorporated HNSCC-specific innervation through injectable 3D human bioengineered trigeminal ganglia, establishing a chip-based innervation–tumor–vasculature tri-interface. Together, this all-human platform captures fundamental determinants of HNSCC progression, including a fully defined ECM, vasculature, and innervation, within a single modular system that is broadly adaptable for interrogating how the tumor microenvironment shapes solid tumor behavior and therapeutic responses.

**Teaser:** HNSCC-on-a-chip integrates defined ECM, vasculature, and innervation to investigate tumor behavior and therapeutic responses.

## INTRODUCTION

The need to study human cancers in their complexity and patient-specificity is driving the generation of innovative, all-human, *in vitro* 3D preclinical models. While *in vivo* models provide the most complete emulation platform for human cancers, they raise significant ethical concerns, and they fall short in replicating human- and patient-specific features (*1*). Among *in vitro* emulation systems, organ-on-a-chip devices are gaining a prominent role in the generation of complex human disease models. These devices are designed to control cell microenvironments and maintain tissue-specific functions, aiming to mimic human physiology (*2*). Early organ-on-a-chip systems connected parallel microchannels separated by a porous membrane to mimic tissue– tissue interfaces. Since then, diverse microfluidic architectures have emerged that achieve similar compartmentalization without membranes, for example, designs that use arrays of micropillars to define central culture chambers or incorporate native biological materials to reproduce natural barriers (*3*). This arrangement recreates tissue-tissue interfaces and closely emulates the minimal functional units found in tissues and organs. The devices allow for real-time, high-resolution imaging of cultured cells and tissues. Moreover, the vascular channels of different organ-specific chips can be linked to study multi-organ physiological coupling (*2–5*). In the context of cancer, organ-on-a-chip platforms allow real-time observation of tumor-immune interactions, drug responses, and cellular behavior under conditions that closely resemble *in vivo* settings. This is particularly valuable for studying cancer cell behavior and the efficacy and limitations of therapeutic interventions in a controlled yet biologically meaningful environment (*2*).

Head and neck squamous cell carcinoma (HNSCC) is the sixth most common cancer worldwide with nearly a million new cases registered in 2020 (*6*). Despite the progress in treatment strategies, advanced HNSCC remains associated with a harsh prognosis (5-year overall survival, <50 %), with a high risk of distant metastasis and local recurrence (*6*). Current therapies for HNSCC include surgery, radiation therapy, chemotherapy, targeted therapy (notably cetuximab, an EGFR inhibitor), and immunotherapy with immune checkpoint inhibitors such as pembrolizumab and nivolumab (*7*). Although these treatments have improved outcomes for some patients, their overall efficacy remains limited, as indicated by the high mortality rate (*8*). Significant barriers still persist in reaching a comprehensive understanding of the biology of these cancers and treatment responses. These limitations primarily arise from the lack of appropriate experimental models that can capture the complexity of HNSCC and its microenvironment (*9*). HNSCC is a complex tissue microenvironment/ecosystem where many different cell types, factors, and extracellular matrix interact (*10*). Key players include cancer-associated fibroblasts (CAFs), vasculature, innervation, and the extracellular matrix (ECM). CAFs stimulate tumor growth, epithelial-to-mesenchymal transition (EMT), invasive potential, and metastasis, and they also modulate the drug sensitivity of cancer cells (*11–13*). Vascularization is fundamental in regulating HNSCC progression: while blood supply provides trophic support to the tumor and a route for metastasis, hypoxia within HNSCC regions distant from vascularization appears to facilitate progression by stimulating radio-resistance (*14, 15*). In recent years, several studies have unraveled the fundamental role of innervation in cancer progression, and in particular, of trigeminal innervation in HNSCC pathogenesis. HNSCC harbors significantly higher neuronal and electrical activity than the corresponding healthy tissue, and innervation actively promotes HNSCC progression and invasiveness (*16, 17*). Furthermore, trigeminal innervation conveys pain, a common and debilitating feature of HNSCC that often appears as one of the first signs of cancer progression (*18*). The extracellular matrix (ECM) itself plays a crucial, yet frequently overlooked role in the progression and treatment resistance of HNSCC. Beyond providing structural support, the ECM actively shapes tumor behavior by influencing cell signaling, migration, and immune evasion (*19*). In HNSCC, aberrant ECM remodeling contributes to a dense, fibrotic tumor microenvironment that can act as a physical and biochemical barrier to therapeutic agents. This is particularly relevant for drug penetration, as the ECM can hinder the uniform distribution of chemotherapeutics and targeted treatments within the tumor mass (*20*). Furthermore, the ECM poses a significant obstacle to the infiltration and efficacy of immune cells and CAR-T cells, which have shown promise in hematologic malignancies but face limited success in solid tumors(*20, 21*).

Recent work has established a small but growing set of HNSCC-on-a-chip and oral-cancer-on-a-chip systems (*9, 22*). Early studies demonstrated that patient-derived HNSCC biopsies could be maintained *ex vivo* on-a-chip for days while exposed to chemotherapeutic agents such as cisplatin and 5-FU, preserving viability and enabling dynamic monitoring of apoptosis and proliferation (*23*). Subsequent platforms extended this approach to radiation testing, showing that perfused tumor slices could reproducibly model dose-dependent cytotoxicity and radiosensitivity in individualized assays (*24, 25*). Other devices refined perfusion design to sustain small tumor fragments for 48h while preserving their histoarchitecture (*26*) or enabled precision-cut HNSCC slices to undergo radiotherapy and cisplatin treatment (*27*). More recently, immunocompetent chips have emerged, such as a humanized 3D microfluidic co-culture combining HNSCC cells, immune cells, and patient serum (*28*). In other works, perfused HNSCC biopsy chips were used to assess transcriptomic and secretome responses after irradiation (*29*), and more recently, oral-cancer on-chip models were proposed to study HNSCC invasion into blood vessels (*30*) and metastasis into the bone (*22*).

Building on these advances, we propose four decisive innovations to address fundamental challenges that remain unresolved in the emulation of the TME. First, we introduce an organ-on-a-chip platform that enables the simultaneous recapitulation of three essential TME components: a three-dimensional patient-derived CAF–HNSCC tumor mass, human trigeminal innervation, and a 3D HNSCC– vasculature interface. Second, we use fully defined engineered extracellular matrices (eECM) with tunable stiffness, composition, and degradability. This represents a critical advance for TME modelling, as existing HNSCC systems, both on-chip and off-chip, either rely on complex yet poorly defined matrices such as Matrigel® or Myogel, or on chemically defined but overly simplified eECMs, including collagen or fibrin. Third, we establish a 3D co-culture system integrating patient-derived HNSCC cells, CAFs, and endothelial cells, thereby partially recapitulating tumor heterogeneity while leveraging the extensive global biobanks of patient-derived HNSCC and CAF lines. Finally, we develop 3D and injectable bioengineered human trigeminal ganglia derived from human embryonic stem cells (hESCs) to model oral cancer–specific innervation.

In a stepwise increase of complexity, we first showed that different eECMs lead to differential self-organization of HNSCC cells, CAFs, and endothelial cells in 3D HNSCC models. We then integrated these 3D HNSCC models into a first organ-on-a-chip device that models the cancer-vasculature interface. We observed ECM-driven differential self-organization of carcinoma cells, collective migration, and vascular invasion. ECM and model complexity strongly influenced the response of 3D HNSCCs to cisplatin, a commonly used chemotherapeutic. We then implemented hESC-derived bioengineered trigeminal ganglia in the complete organ-on-a-chip device that models innervation/cancer and cancer/vascular interfaces. We observed rapid and abundant innervation of HNSCC/CAFs, and innervation-dependent radical increase in tumor size, accompanied by altered collective migration dynamics. Taken together, our study proposes an innovative innervated and vascularized 3D model of HNSCC, developed in completely defined extracellular matrices with tunable stiffness, composition, and degradability, which allows fine control of the tumor microenvironment and permits the study of HNSCC behavior at the tumor-vasculature and tumor-innervation interfaces. The platform can be further easily adapted to the investigation of virtually any biological question in which controlled and defined 3D eECM, innervation, and vasculature are needed in an *in vitro* or *ex vivo* setting.

## RESULTS

### 1. Fabrication of an organ-on-a-chip device for the emulation of the interface between HNSCC, vasculature, and trigeminal innervation

We generated an organ-on-a-chip device that allows the emulation of the 3D interface between HNSCC tumor mass, vasculature, and trigeminal innervation (Figure 1A). The device is composed of 4 adjacent, sequential chambers (Figure 1B): in the middle, two chambers for the 3D culture of HNSCC and human bioengineered trigeminal ganglia (hbTG), on the right a vascular chamber, on the left a channel to supply medium to the hbTG (Figure 1B). The 3D HNSCC chamber is separated from the adjacent vascular chamber by micropillars (Figure 1C). These micropillars allow containment of the hydrogel-cells mixture (3D HNSCC) during the crosslinking phase. The pillars are discontinuous to permit direct contact and formation of a continuous endothelial barrier between the 3D HNSCC and the vascular chamber upon seeding of endothelial cells in the latter. Culture medium for the 3D HNSCC is supplied via the vascular chamber. The 3D HNSCC chamber is connected to the 3D trigeminal chamber via microgrooves (5 μm heigh, 10 μm wide, 150 μm long, with a 50 μm spacing between them; Figure 1C) that allow separation of media between the two compartments and the passage of axons, but not of whole cells. This set up aims to reproduce the anatomical separation between the soma of trigeminal neurons, located in the trigeminal ganglion, and the innervation of HNSCC that occurs in the oral cavity. The 3D trigeminal ganglia are supplied with culture medium via a dedicated medium channel, located on the extreme left side of the chip (Figure 1B). We further produced a simplified version of the device that includes only the 3D HNSCC and the vascular chamber, for experiments focused on the tumor-vasculature interface without inclusion of trigeminal innervation (Figure 1B). To characterize molecular transport across the different compartments of the microfluidic devices, we performed diffusion assays using fluorescent tracers with distinct molecular weights, namely Rhodamine B (500 Da) and dextran (2000 kDa) (Figure 1D). In the reduced device design, when fluorescent tracers were introduced into the vascular channel, Rhodamine rapidly diffused across the micropillar interface into the adjacent 3D HNSCC chamber, with detectable fluorescence observed within 5 minutes and a progressive increase in the tumor compartment over time. In contrast, dextran diffusion from the vascular channel into the tumor chamber was not observed even after 1 hour, with fluorescence remaining confined to the vascular channel (Figure 1D, top panels). We next assessed diffusion between the trigeminal ganglion chamber and the 3D HNSCC compartment through the connecting microchannels in the complete design. Upon introduction of fluorescent tracers into the trigeminal medium channel, Rhodamine readily diffused into the trigeminal compartment (within 5 minutes) and partially through the microchannels into the tumor chamber within 1 hour (Figure 1D, bottom panels). Conversely, dextran diffusion was strongly limited, with fluorescence largely restricted to the trigeminal medium channel, with minimal signal detected in the trigeminal ganglion compartment, and no detectable signal in the tumor compartment after 1 hour.

**Figure 1.**
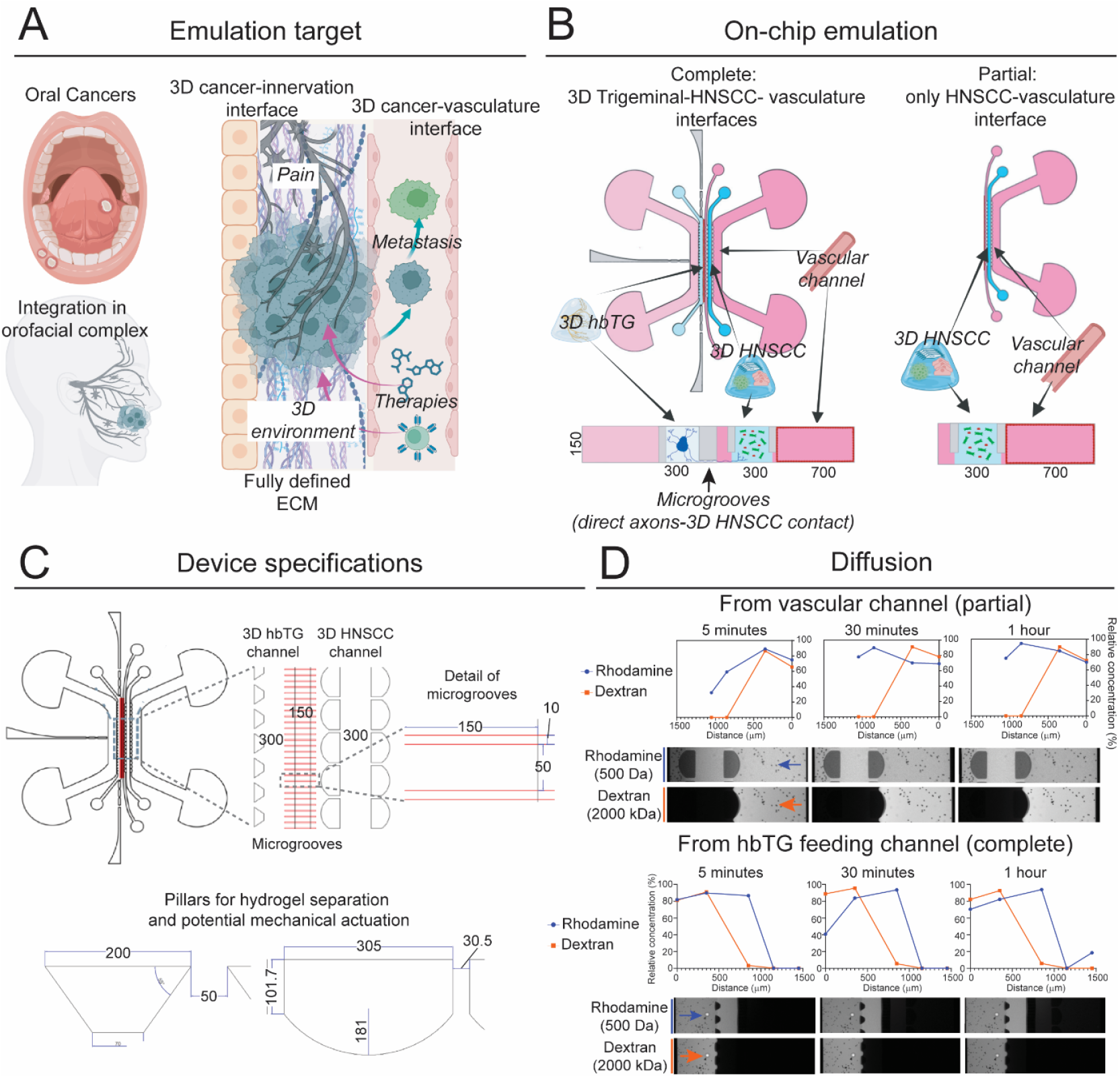
Organ-on-a-chip device for the emulation of the interface between HNSCC, vasculature, and trigeminal innervation. **A)** Schematic representation of the emulation target of the HNSCC-on-a-chip device. **B)** Basic design of the Complete HNSCC-on-a-chip device (left) and Reduced HNSCC-on-a-chip device (right). **C)** Details of the device design. All dimensions are expressed in μm. **D)** Diffusion characterization of the microfluidic devices using fluorescent tracers (i.e. Rhodamine B (500 Da) and Dextran (2000 kDa)). Top panels show the diffusion of rhodamine B (blue line) and dextran (red line) from the vascular channel into the adjacent 3D HNSCC compartment in the partial device, while bottom panels show the diffusion from the trigeminal medium channel into the trigeminal ganglion and 3D HNSCC compartments in the complete device. Representative fluorescence microscopy images and corresponding percentage of the relative concentration profiles are shown for Rhodamine B and dextran at 5 minutes, 30 minutes, and 1 hour. Blue and red arrows indicate the diffusion direction of rhodamine and dextran, respectively.

### 2. The composition of the engineered extracellular matrix (eECM) induces rapid and differential self-organization patterns of co-cultured HNSCCs, CAFs, and endothelial cells

We first assessed the effect of different engineered extracellular matrices (eECMs) on the behavior of HNSCC cells (HNSCCs), cancer associated fibroblasts (CAFs), and endothelial cells (HUVECs), collectively referred to as HCHs, in isolated 3D dome cultures (Figure 2A). We focused on fully defined eECMs, as our aim is to create a model in which the eECM has defined composition, desired mechanical properties, and controlled proteolytic degradation. For this purpose, we generated a fully defined, complex eECM with the above mentioned properties by modifying hyaluronan with bicyclo[6.1.0]non-4-yn-9-ylmethyloxycarbonyl]-1,8-diamino-3,6-dioxaoctane (HA-BCN), allowing for cross-linking using azide-terminated peptides and linear and multi-arm poly(ethylene glycol) (PEG-Azn) by strain-promoted alkyne–azide cycloaddition (SPAAC). We utilized 8-arm PEG-Azide (PEG-Az8) and a protease degradable peptide cross-linker with terminal azide moieties (VPM-Az2). We further included collagen I and recombinant human laminin by entrapment (HA/PEG/VPM/ColI/Lam: HPVCL; Figure 2B). Cross-linking of the hydrogel commences immediately upon mixing HA-BCN with the cross-linkers and the hydrogels reach their final stiffness after approximately 1 hour. We compared this eECM with a non-protease degradable version (where we omitted VPM, HA/PEG/ColI/Lam: HPCL) and with a defined, natural, and patho-physiologically relevant eECM widely used in preclinical models of human cancers: fibrin (Figure 2B). We mixed HNSCC/CAFs/HUVECs (HCH) at a 1:3:1 ratio and co-cultured them in HPVCL, HPCL, or Fibrin, at a total concentration of 15^*^10^6^ cells/mL (Figure 2C). HCHs cultured in fibrin remained spread throughout the eECM, while HUVECs formed analogous vascular networks integrated within the tumor mass. HNSCCs and CAFs self-organized into large separate domains, with HNSCCs occupying the center of the eECM and CAFs distributed on the periphery (Figure 2D). CAFs also tended to migrate out of the eECM onto the plastic surrounding the dome (Figure 2D; arrowheads). HCHs cultured in HPVCL showed a rapid and robust migration towards the center of the eECM, followed by self-organization into discrete, highly dense tumor masses within the first 24 hours. This organization was maintained grossly unaltered for at least 4 days (Figure 2E). Within these masses, HUVECs rapidly gave rise to ramified vascular structures in direct contact with HNSCC and CAFs. HNSCC and CAFs distributed without large scale zonation with the tumor mass (Figure 2E). Notably, this self-organization did not require proteolytic activity, as we observed the same collective cells behavior in HPCL eECMs, which are devoid of the protease-degradable VPM crosslinker (Figure 2F).

**Figure 2.**
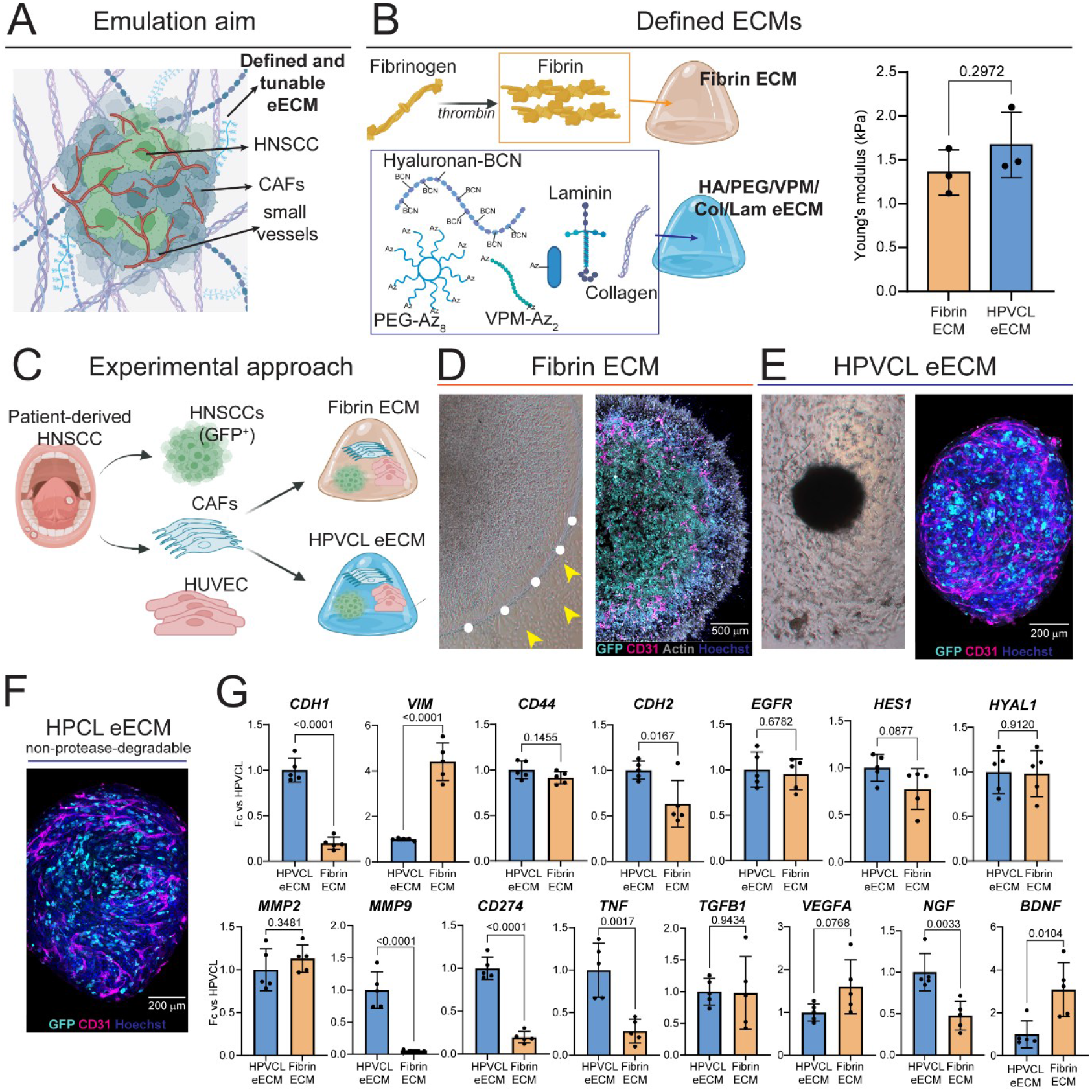
Emulating the HNSCC/ECM interface: co-culture of HNSCC cells, cancer-associated fibroblasts (CAFs), and endothelial cells (HUVEC) in different fully defined extracellular matrices (ECM). **A)** Schematic representation of the tumor microenvironment features emulated by the system. **B)** Composition of Fibrin ECM and Hyaluronic acid-BCN, PEG-Az8, Collagen, Laminin, VPM-Az2 (HPVCL) eECM. Bar plot: comparison of Young’s Modulus of Fibrin ECM and HPVCL eECM, as determined by nanoindentation. **C)** Experimental approach for co-culturing HNSCC cells, CAFs, and HUVECs (HCHs) in Fibrin ECM and HPVCL eECM. **D)** HCHs organization after 4 days of culture in Fibrin ECM. Left: brightfield. HCHs are sparse throughout the ECM and cells also invade the plastic dish surrounding the ECM dome (white dots: hydrogel border. Arrowheads: cells leaving the hydrogel). Right: maximum intensity projection of 3D immunofluorescent staining showing the relative positioning of HNSCC cells (GFP^+^, in cyan), CAFs (only stained via an actin marker, in white) and HUVEC (C31^+^, in magenta). HNSCCs occupy the center of the 3D construct, while CAFs surround them. HUVECs form sparse capillary networks. Blue: Hoechst3342 (nuclei). N=4 independent replicates. **E)** HCHs organization after 4 days of culture in HPVCL. Left: brightfield image showing HCHs self-organize in a highly dense tumor mass located at the center of the eECM. Right: maximum intensity projection of immunofluorescent staining showing the relative positioning of HNSCC cells (GFP^+^, in cyan), CAFs (only stained via an actin marker, in white) and HUVEC (C31^+^, in magenta). Blue: Hoechst3342 (nuclei). HUVEC cells form dense capillary networks, adjacent to highly dense HNSCC masses surrounded by CAFs. N=4 independent replicates. **F)** Maximum intensity projection of immunofluorescent staining showing the relative positioning of HNSCC cells (GFP^+^, in cyan), CAFs (only stained via an actin marker, in white) and HUVEC (C31^+^, in magenta) in HPCL eECM, in which the protease degradable VPM-Az2 crosslinker has been omitted, making the eECM degradable only by hyaluronidases. Blue: Hoechst3342 (nuclei). N=4 independent replicates. **G)** Real time quantitative PCR analysis of changes in gene expression between HCHs cultured in Fibrin ECM or HPVCL eECM for 4 days. N = 5 independent replicates per condition. T-test, exact p values are reported onto each comparison. Abbreviations – CAFs; cancer associated fibroblasts; ECM: extracellular matrix; eECM: engineered extracellular matrix; HNSCCs: human head and neck squamous cell carcinomas; HUVECs: human umbilical vein endothelial cells.

We then compared the effects of culturing HCHs for 4 days in two different ECMs on the bulk expression of genes involved in HNSCC behavior, ECM remodeling, and HNSCC-TME interactions by real time quantitative PCR (RT-qPCR). HCHs cultured in fibrin or HPVCL showed significant differences in the expression of markers of epithelial-mesenchymal identity and transition, with fibrin promoting the expression of *VIM* (coding for Vimentin, mesenchymal marker) and HPCVL the expression of *CDH1* and *CDH2. CD44*, marker of cancer stemness and receptor for hyaluronic acid (*31*–*33*), the HNSCC driver *EGFR* (*34*), and the Notch pathway effector *HES1* showed comparable expression in HCHs cultured in the two ECMs. Culture in HPCLV induced on the other hand 10-fold higher expression of *MMP9* (matrix metalloprotease 9; Figure 2G), protease involved in ECM remodeling associated with HNSCC invasion (*35*). The expression of *MMP2*, another protease involved in ECM remodeling, and of *HYAL1*, responsible for hyaluronic acid degradation in HNSCC progression (*36, 37*), were unaffected by the ECM composition (Figure 2G). HPVCL further induced higher expression of key regulators of HNSCC immune escape, such as *CD274* (PD-1L) and *TNF* (*38*) (Figure 2G). Genes involved in tumor innervation were also affected by the ECM composition, with *NGF* being more expressed by HCHs cultured in HPVCL, and *BDNF* more expressed by HCHs cultured in fibrin (Figure 2G). Taken together, these data indicate that the choice of ECM exerts significant effects on HNSCC self-organization and gene expression.

### 3. eECM composition induces differential self-organization, migration, and vasculature invasion of HNSCC at the tumor-vasculature interface

We increased the complexity of our modeling of the TME by integrating the fibrin-based and HPVCL-based 3D HNSCCs described in section 2 into our partial HNSCC-on-a-chip device, which allows the emulation of a 3D HNSCC-vascular interface (Figure 3). In this system, we first investigated the effects of eECM composition on the behavior of 3D HCHs and tumor cells migration towards vasculature, a common route for metastasis (*39*) (Figure 3A). To this end, we resuspended HCHs either in HPVCL-eECM or fibrin-ECM, injected them in their dedicated chamber adjacent to the HUVEC-coated vascular channel, and monitored their behavior over time (Figure 3B). Already after 24 hours, cells displayed strong eECM-specific self-organization. While HCHs cultured in fibrin-eECM showed an elongated morphology and were distributed evenly across the matrix (Figure 3C), HCHs cultured in HPVCL-eECM self-organized in multiple dense tumor masses (Figure 3D). Within two days, HCHs cultured in fibrin-eECM migrated over multiple invasion fronts towards the vascular compartment and invaded it along the entire tumor-vasculature interface (Figure 3C-C’’). HCHs cultured in HPVCL-eECM showed a radically different behavior: merged in tumor masses, they formed discrete invasion fronts that migrated as a collective unit through the eECM towards the vascular chamber, which they invaded by day three (Figure 3D-D’’). We performed immunofluorescent staining followed by confocal microscopy to investigate the spatial relationship between the different cell types that compose the 3D HNSCC and their dynamic during tumor extravasation. We observed that, in HPVCL-eECM, migration through the eECM and invasion of the vascular channel was driven by CAFs, which showed higher degrees of proliferation and created a path for successive HNSCC migration (Figure 3E). By day 4, we observed multiple discrete invasion fronts into the vasculature, led by CAFs and followed by HNSCCs (Figure 3E’-E’’’). Overall, the vascular compartment exerted a strong attractive effect on the tumor mass, which was not limited to vascular invasion: in several instances, we observed CAFs migrating around and below the endothelial cells, engulfing the vessel (Figure 3F-F’’’). By day 7, most migration fronts invaded the vessel, causing the loss of the endothelial barrier between the vascular and the 3D HNSCC. However, not all HCHs invaded the vessel: we observed the persistence of non-proliferating, yet alive HCH-masses within the HPVCL-eECM well after the invasive fronts entered the vasculature (Figure 3G-G’’). HCHs cultured in fibrin-eECM showed a radically different behavior, as they indiscriminately invaded the vascular chamber by day 7, creating a continuum with those remaining within the eECM. In this case, CAFs and HNSCC formed multiple, continuous invasion routes from the eECM to the vasculature, with no discernible morphological distinction (Figure 3H-H’’’). Taken together, these results show that the composition of the eECM radically shapes the proliferation, migration, and self-organization of HNSCC-derived cancer cells and fibroblasts, and the dynamic of their extravasation.

**Figure 3.**
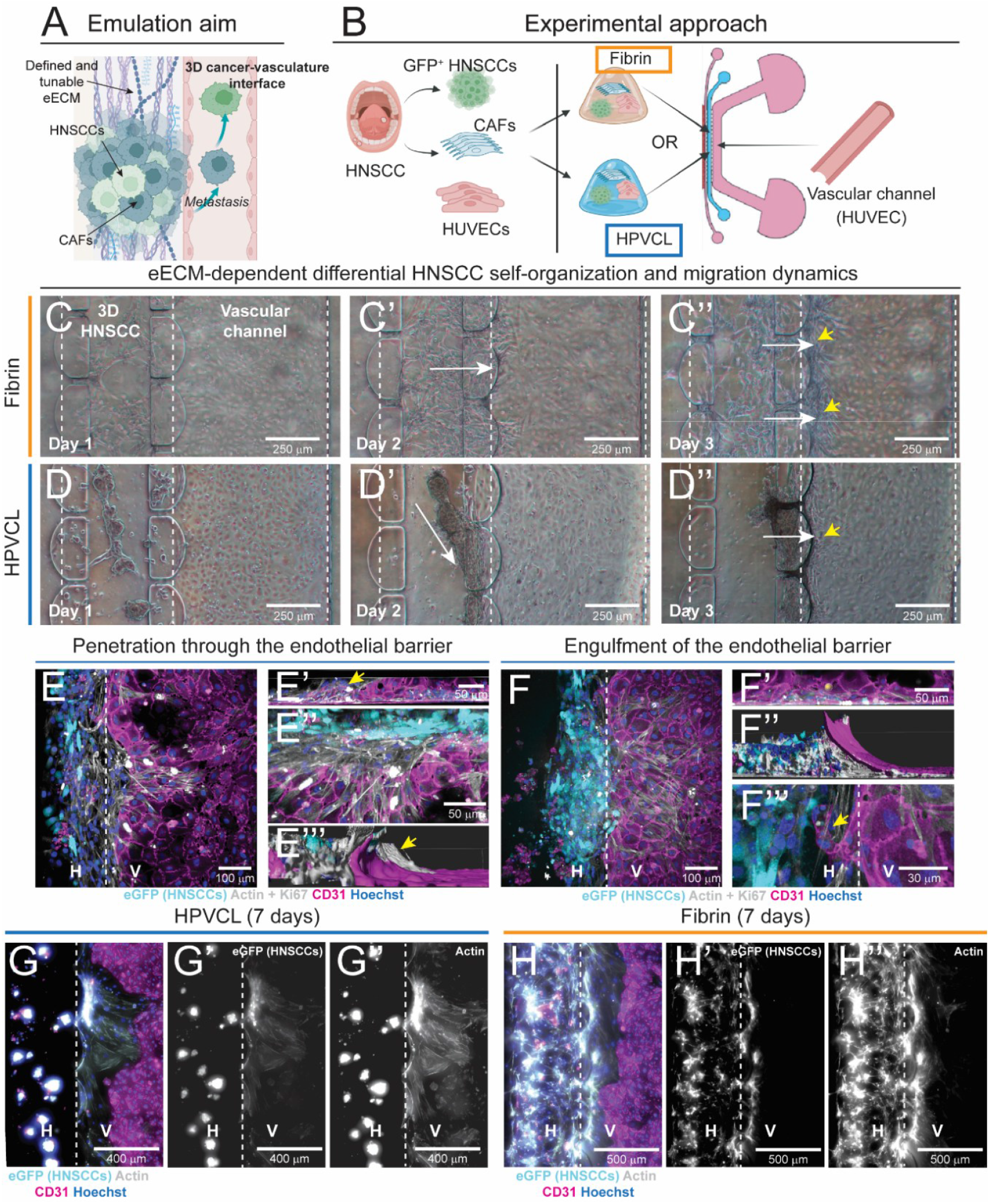
Emulation of the 3D HNSCC/ECM/vascular interface. **A)** Schematic representation of the emulation aim. **B)** Experimental approach to co-culture HNSCCs, CAFs, and HUVEC (together referred to as HCHs) in different ECMs at the interface with a HUVEC-lined vascular channel. N = 3 per condition, per time point (day 4 and day 7). **C-C’’)**Brightfield images showing the organization, migration, and vascular invasion pattern of HCHs when cultured in fibrin ECM. White arrows indicate the migration direction; yellow arrowheads indicate some of the areas of invasion of the vascular channel. **D-D’’)** Brightfield images showing the organization, migration, and vascular invasion pattern of HCHs when cultured in HPVCL eECM. White arrows indicate the migration direction; yellow arrowheads indicate some of the areas of invasion of the vascular channel. **E-E’’’)** Immunofluorescent staining showing HCHs cultured in HPVCL eECM invading the vascular channel through the endothelial barrier, day 4. Cyan: GFP (HNSCCs); white: actin (marking strongly CAFs) + Ki67; Magenta: CD31 (HUVECs); blue: Hoechst3342 (nuclei). H = HNSCC channel; V = vascular channel. **E**: top view of 3D confocal imaging, showing a top view of the invasion front. **E’**: view of the endothelial barrier and the invasion front from inside the vascular channel. The continuous endothelial barrier is interrupted by HCHs (yellow arrowhead). **E’’**: detail of the invasion front, showing CAFs leading the front, followed by HNSCCs. **E’’’**: 3D rendering on the vascular invasion front (yellow arrowhead, side view. **F-F’’’)** Immunofluorescent staining showing HCHs cultured in HPVCL eECM engulfing the vascular channel, day 4. Cyan: GFP (HNSCCs); white: actin (marking strongly CAFs) + Ki67; Magenta: CD31 (HUVECs); blue: Hoechst3342 (nuclei). H = HNSCC channel; V = vascular channel. **F**: top-view, showing apparent invasion of the vascular channel. Yet, observation from within the vascular channel (**F’**) does not show any HCHs. 3D rendering (**F’’**) highlights migration of HCHs, and mostly CAFs, below the endothelial layer, engulfing the vascular chamber. **F’’**: evidence of either anastomosis or vascular sprouting from the main vascular channel into the 3D HNSCC compartment. **G-G’’**) Day 7 of culture of HCHs in HPVCL eECM adjacent to vascular channels. **G**: maximum projection intensity, top-view, Cyan: GFP (HNSCCs); white: actin (marking strongly CAFs) + Ki67; Magenta: CD31 (HUVECs); blue: Hoechst3342 (nuclei). H = HNSCC channel; V = vascular channel. **G’**: only GFP channel; **G’’**: only actin channel. **H-H’’**) Day 7 of culture of HCHs in fibrin ECM adjacent to vascular channels. H: maximum projection intensity, top-view, Cyan: GFP (HNSCCs); white: actin (marking strongly CAFs) + Ki67; Magenta: CD31 (HUVECs); blue: Hoechst3342 (nuclei). H = HNSCC channel; V = vascular channel. **H’:** only GFP channel; **H’’**: only actin channel.

### 4. ECM composition and model complexity influence HNSCC response to chemotherapy

We further investigated whether the eECM composition affects the response of 3D HNSCC models to cisplatin, a chemotherapeutic commonly used to treat HNSCC (*40*). To this end, we cultured HCHs either in Fibrin or in HPVCL as described above. After 24 hours, we supplemented the culture medium with 3 μg/mL cisplatin and analyzed the 3D HNSCCs after 72 hours of treatment (Figure 4A). In response to cisplatin, HCHs cultured in fibrin formed a dense tumor core in which cells survived 3 days of treatment, with vascular structures severely damaged, yet still detectable (Figure 4B-B’). Cells located at the periphery of the ECM were eradicated by cisplatin as indicated by a cell-free halo at the ECM border (Figure 4B), and by the accumulation of cleaved caspase 3 at the periphery of the tumor area (Figure 4B’’). Notably, no cells were detected on the plastic surrounding the fibrin ECM (compare Figure 4A with Figure 2D), indicating that cisplatin efficiently killed cells out of the ECM, at its periphery, or prevented their migration beyond the fibrin scaffold. In contrast, HCHs grown in HPVCL showed a strikingly higher sensitivity to cisplatin, as the self-assembled tumor masses were completely lost, with only a few scattered round cells detected in the eECM (Figure 4C-C’). Most of the few visible cells showed cleaved caspase 3 expression, indicative of ongoing apoptosis (Figure 4C’’).

**Figure 4.**
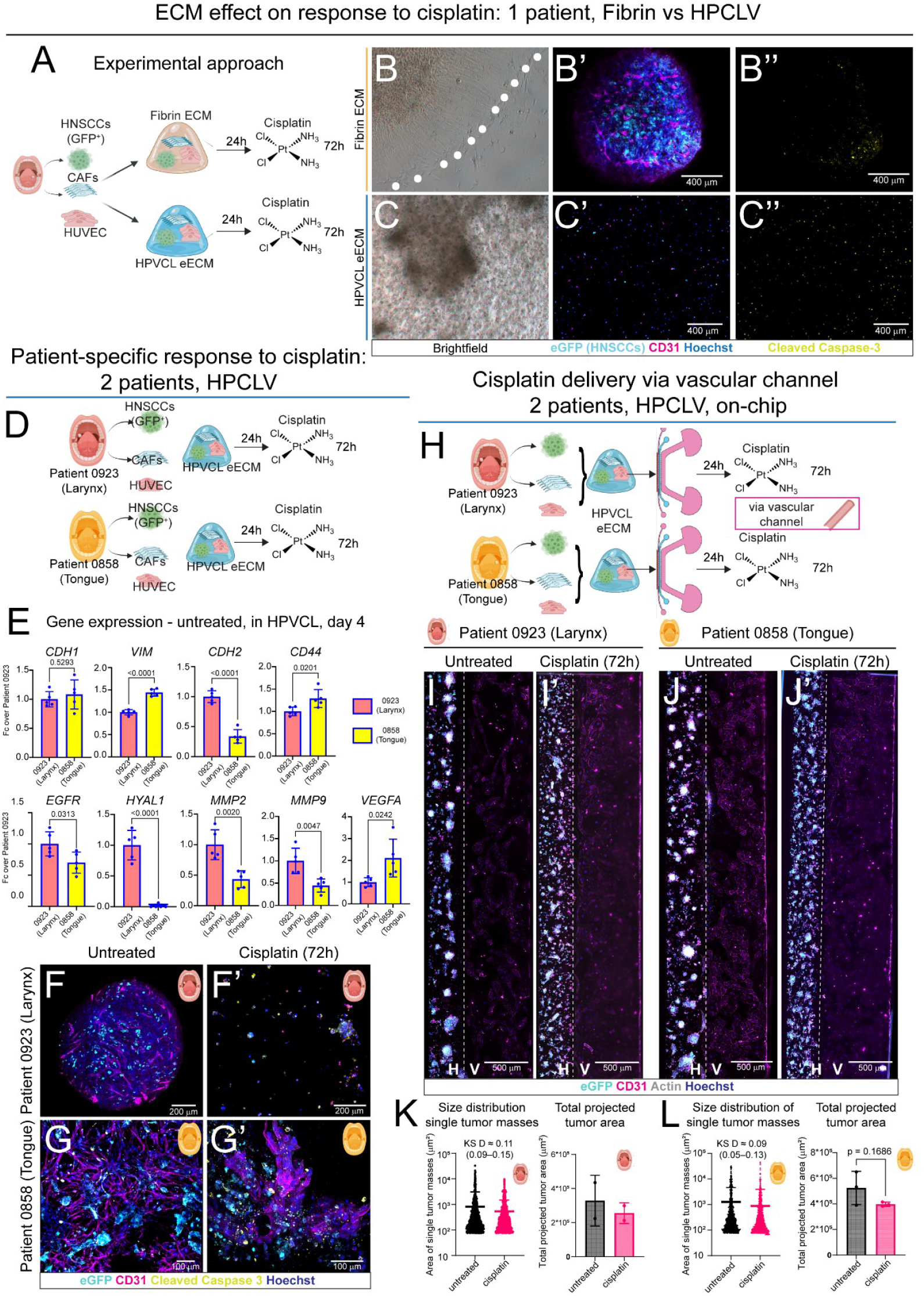
(next page). Effect of ECM, patient variability, model complexity and delivery route on response to cisplatin treatment. **A)** Experimental approach: effect of ECM composition on response to cisplatin. **B-B’’)** Brightfield image (B) and immunofluorescent staining (maximum intensity projection, B’-B’’) of HNSCC, CAFs and HUVECs (HCH) cultured in fibrin and treated with cisplatin (3μg/mL) from 24 hours of culture and for 4 days. N= 4 independent replicates per condition. **B**, white dots: limit of fibrin ECM. **B’**: cyan – GFP/HNSCC; magenta – CD31 (HUVEC); blue – Hoechst3342 (nuclei). **B’’**: yellow – cleaved caspase 3 (apoptosis). **C-C’’)** Brightfield image (C) and immunofluorescent staining (maximum intensity projection, C’-C’’) of HNSCC, CAFs and HUVECs (HCH) cultured in HPVCL and treated with cisplatin (3μg/mL) from 24 hours of culture and for 3 days. N= 4 independent replicates per condition. **C’**: cyan – GFP/HNSCC; magenta – CD31 (HUVEC); blue – Hoechst3342 (nuclei). **C’’**: yellow – cleaved caspase 3 (apoptosis). **D)** Experimental approach: patient-specific response to cisplatin in the same HPVCL eECM. **E)** Quantitative real time PCR analysis of differential expression of selected genes between 0858-HNSCC/CAFs/HUVECs and 0923-HNSCC/CAFs/HUVECs cultured in the same HPVCL eECM for 4 days. N = 5 per condition; t-test: exact p-value indicated on each comparison. **F-F’)** Immunofluorescent staining (maximum intensity projection of 3D stack) showing 0923-HNSCC/CAFs/HUVECs cultured in HPVCL in control conditions (F) and after treatment with cisplatin (3μg/mL) from 24 hours of culture and for 3 days (F’). Cyan: GFP/HNSCC; magenta: CD31 (HUVEC); blue: Hoechst3342 (nuclei); yellow: cleaved caspase 3 (apoptosis). N= 4 independent replicates per condition. **G-G’)** Immunofluorescent staining (maximum intensity projection of 3D stack) showing 0858-HNSCC/CAFs/HUVECs cultured in HPVCL in control conditions (G) and after treatment with cisplatin (3μg/mL) from 24 hours of culture and for 3 days (G’). Cyan: GFP/HNSCC; magenta: CD31 (HUVEC); blue: Hoechst3342 (nuclei); yellow: cleaved caspase 3 (apoptosis). N= 4 independent replicates per condition. **H)** Experimental approach: culture of 0858-HNSCC/CAFs/HUVECs and 0923-HNSCC/CAFs/HUVECs in HPVCL in contact with vascular channel, and treatment with cisplatin via the vascular channel. **I)** Immunofluorescent staining (maximum intensity projection of 3D stack) of 0923-HNSCC/CAFs/HUVECs in HPVCL cultured in control conditions for 4 days. H = HNSCC channel; V = vascular channel. Cyan: GFP/HNSCC; magenta: CD31 (HUVEC); blue: Hoechst3342 (nuclei); white: actin (GFP^+^Actin^+^ = HNSCC; GFP^-^ Actin^+^ = CAFs; CD31^+^Actin^+^ = HUVECs). N= 3 independent replicates. **I’)** Immunofluorescent staining (maximum intensity projection of 3D stack) of 0923-HNSCC/CAFs/HUVECs in HPVCL cultured with cisplatin (3μg/mL) from 24 hours of culture and for 3 days. H = HNSCC channel; V = vascular channel. Cyan: GFP/HNSCC; magenta: CD31 (HUVEC); blue: Hoechst3342 (nuclei); white: actin (GFP^+^Actin^+^ = HNSCC; GFP^-^ Actin^+^ = CAFs; CD31^+^Actin^+^ = HUVECs). N= 3 independent replicates. **J)** Immunofluorescent staining (maximum intensity projection of 3D stack) of 0858-HNSCC/CAFs/HUVECs in HPVCL cultured in control conditions for 4 days. H = HNSCC channel; V = vascular channel. Cyan: GFP/HNSCC; magenta: CD31 (HUVEC); blue: Hoechst3342 (nuclei); white: actin (GFP^+^Actin^+^ = HNSCC; GFP^-^ Actin^+^ = CAFs; CD31^+^Actin^+^ = HUVECs). N= 3 independent replicates. **J’)** Immunofluorescent staining (maximum intensity projection of 3D stack) of 0923-HNSCC/CAFs/HUVECs in HPVCL cultured with cisplatin (3μg/mL) from 24 hours of culture and for 3 days. H = HNSCC channel; V = vascular channel. Cyan: GFP/HNSCC; magenta: CD31 (HUVEC); blue: Hoechst3342 (nuclei); white: actin (GFP^+^Actin^+^ = HNSCC; GFP^-^ Actin^+^ = CAFs; CD31^+^Actin^+^ = HUVECs). N= 3 independent replicates. **K, L)** Distribution of projected area of each single tumor mass (left) and of total tumor area per chip (right) detected in 0923-HNSCC/CAFs/HUVECs-vascular chips (K) and 0858-HNSCC/CAFs/HUVECs-vascular chips (L). Cisplatin treatment causes a strong decrease in the number of larger tumor masses but only a minor decrease in the overall tumor area. Distribution analysis: chip-level Kolmogorov–Smirnov analysis, KSD = KS distance. Comparison of total tumor area: T-test, exact p value is indicated on the graph.

We then investigated whether this strong sensitivity to cisplatin in HPVCL was specific to the HNSCC line used, line 0923 (isolated from a laryngeal HNSCC, (28)). We thus performed the same experiment using another line from our cell bank, line 0858 (isolated from a tongue HNSCC; figure 4D). We first cultured 0923-HNSCCs and 0858-HNSCCs with the same CAFs and HUVECs, and in the same HPVCL eECM, to investigate patient-specific changes in HCHs self-organization. 0858-HNSCCs/CAFs/HUVECs still formed a mass in the center of the eECM, but much less dense, with a higher number of capillaries, and higher spacing between HNSCC and CAFs (Figure 4F, G). The lower tumor mass density in the eECM was associated with an overall lower expression of genes coding for ECM remodelers: 0858-HCHs expressed significant lower levels of *MMP2, MMP9*, and *HYAL1*, all involved in the degradation of VPM-Az2 or hyaluronic acid, necessary to remodel HPVCL. 0858-HCHs expressed overall slightly higher levels of *VIM, CD44*, and *VEGF*, and lower levels of *EGFR* and *CDH2*. The two models expressed similar levels of *CDH1* and *CD44*. We treated both 0858-HCHs and 0923-HCHs with cisplatin for 3 days and analyzed their response via immunofluorescent staining followed by 3D confocal microscopy. Line 0923 confirmed its high sensitivity to cisplatin, with only single, round cells detectable in the construct and widespread cleaved caspase 3 signal (figure 4F’). 0858-HCHs showed a less severe response. While the capillary network was strongly affected and cells died in response to the treatment, HCH masses could still be observed within the eECM (Figure 4G’).

We then set out to increase the pathophysiological relevance of our model by culturing 3D 0858-HCHs and 0923-HCHs in HPVCL, adjacent to a vascular channel on chip, as described above (Figure 3A), and by delivering cisplatin via the vascular channel, thereby more closely resembling clinical systemic delivery (Figure 4H). Thus, we cultured the two 3D HNSCC combinations on chip for 4 days, providing culture medium via the vascular channel. A subset of chips was treated with cisplatin supplied via the vascular channel, beginning after 24 hours and renewed daily until day 4 (Figure 4H). Notably, on-chip, untreated 0858-HCHs and 0923-HCHs showed highly similar self-organization patterns, both displaying organization into discrete tumor masses within the 3D HNSCC chamber (Figure 4I and 4J; day 4). 0858-HCHs and 0923-HCHs treated with cisplatin showed a much more similar response in this system that what observed when cultured in isolated 3D eECMs. After 3 days of cisplatin treatment, in both systems we observed a moderate but consistent reduction in the size of the individual tumor masses (0923 KS D ≈ 0.09 [0.05-0.13], and 0858 KS D ≈ 0.11 [0.09-0.15]), with the decrease of larger tumors (Figure 4K, L) and the formation of multiple smaller tumors. Yet, the total tumor mass did not significantly decrease within 3 days, either for patient 0923 or for patient 0858 (Figure 4K, L). Taken together, these data highlight the importance of the choice of ECM, model complexity, and delivery route to investigate the effects of chemotherapy on complex human tumors.

### 5. Generation of 3D hESC-derived human bioengineered trigeminal ganglia (hbTG)

We then set out to generate 3D human bioengineered trigeminal ganglia (hbTG) in a fully defined and injectable 3D eECM. To this end, we adapted an existing protocol that generates diverse populations of trigeminal neurons, including neurons responsive to heat, cold, and inflammatory pain, as well as trigeminal glial cells from human pluripotent stem cells (Figure 5A) (*41*). We first differentiated human embryonic stem cells (hESCs) into a mixed population of trigeminal neural crest and placode cells in a classic 2D culture. We then detached the trigeminal precursors obtained, encapsulated them in an eECM composed of HA-BCN, Peg-Az8, cRGD, and Laminin 521 (HPRL), and cultured them in neuronal differentiation medium for up to 40 days (Figure 5A). Via this approach we obtained organization of trigeminal precursors into dense trigeminal ganglia-like structures that sprouted bundled axons into the eECM (Figure 5B-B’’). At the border of the eECM, axons showed a stronger tendency to bundle into nerve-like structures (Figure 5D, E). Notably, we detected SOX10-expressing glial lineage cells interspersed among the neuronal soma within the ganglia as well as along the axons projected into and outside the eECM (Figure 5D, E). hESC-derived bioengineered trigeminal ganglia (hbTG) expressed the trigeminal markers *PAX3* and *SIX1*, as well as the neuroblasts marker *BRN3A* (Figure 5F). Expression of *PAX3* and *SIX1* was highest at the trigeminal precursor stage to then decrease during neuronal differentiation, in accordance with their expression profile during trigeminal development *in vivo* (*42*–*44*). Neuronal differentiation was accompanied by a strong increase in the expression of *CALCA* (marker for nociceptive neurons) (*45*) and of *SCN10A*, coding for the voltage-gated sodium channel Nav1.8 responsible for the transmission of action potentials (*46*). We further detected increased expression of *PIEZO2* (marker for sensory neurons responsive to mechanical stimulation), *TRPM8* (marker for cold-responsive sensory neurons), and to a lesser extent *TRPV1* (marker for heat-responsive neurons) (*47, 48*). The strong expression of nociceptive markers and *SCN10A* was accompanied by action potential transmission in response to stimulation with ATP (nociceptive signal; supplementary Video S1).

**Figure 5.**
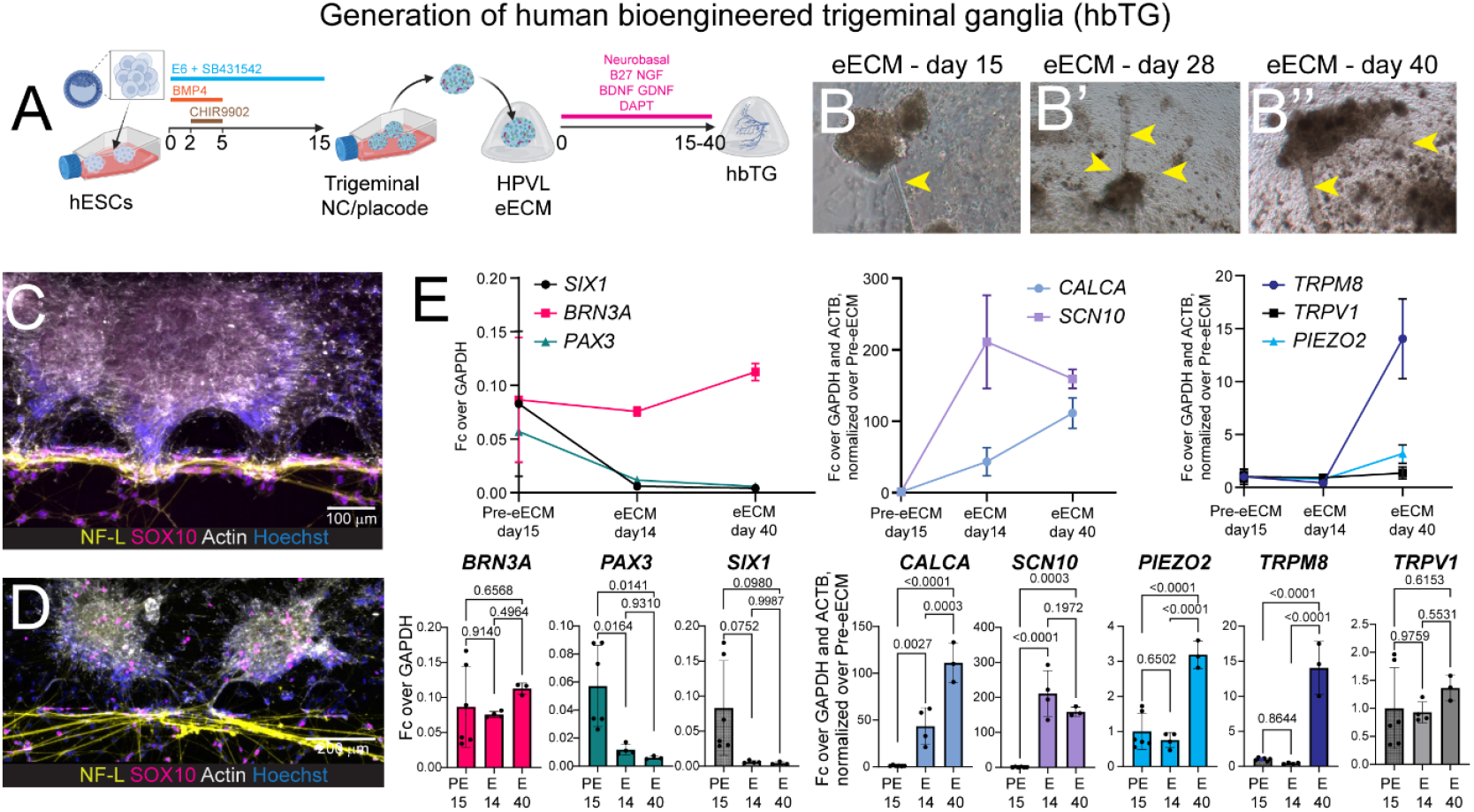
Generation of 3D hESC-derived bioengineered trigeminal ganglia (hbTG). **A)** Experimental approach. **B-B’’)** Brightfield images showing hbTG at different stages of differentiation. Yellow arrowheads show bundled axons. **C, D)** Immunofluorescent staining (slice sum from 3D stack) of hbTG cultured in HPVL for 7 days on chip. Yellow: neurofilament light; red: SOX10; white: actin; blue: Hoecsht3342. N= 3 independent replicates. **E)** Quantitative real time PCR analysis of the expression of markers of trigeminal identity and different trigeminal neurons subpopulations in hbTG at different stages of differentiation. PreeECM day 15 (PE15): n = 6; eECM day 14 (E14): n = 4; eECM day 40 (E40): n = 3. One-way ANOVA with Tukey correction for multiple comparisons; exact p value indicated on each comparison.

### 6. Trigeminal innervation induces increased tumor growth but reduced extravasation of HNSCCs/CAFs

We assessed the effect of trigeminal innervation on the behavior of HCHs cultured in HPVCL on-chip. To this end, we injected 3D hbTGs and 0923-HCHs into the central adjacent chambers connected by microchannels, to allow axon projection from the hbTG to the HNSCC chamber. We further generated a vascular channel in the right-most channel, in direct contact with the HNSCC compartment (Figure 6A). All tumor structures in hbTG-containing chips were profusely innervated. Axons followed the borders of the tumor masses (Figure 6C-D’’) and preferentially grew along the tumor surface rather than within the eECM. Axons did not show preferential interaction with CAFs or HNSCCcs, innervating both. Notably, innervation was very abundant at many tumor-vascular interfaces, creating complex neuro-vascular/tumor bundles (Figure 6D) that followed or guided HNSCC invasion of the vascular channel.

**Figure 6.**
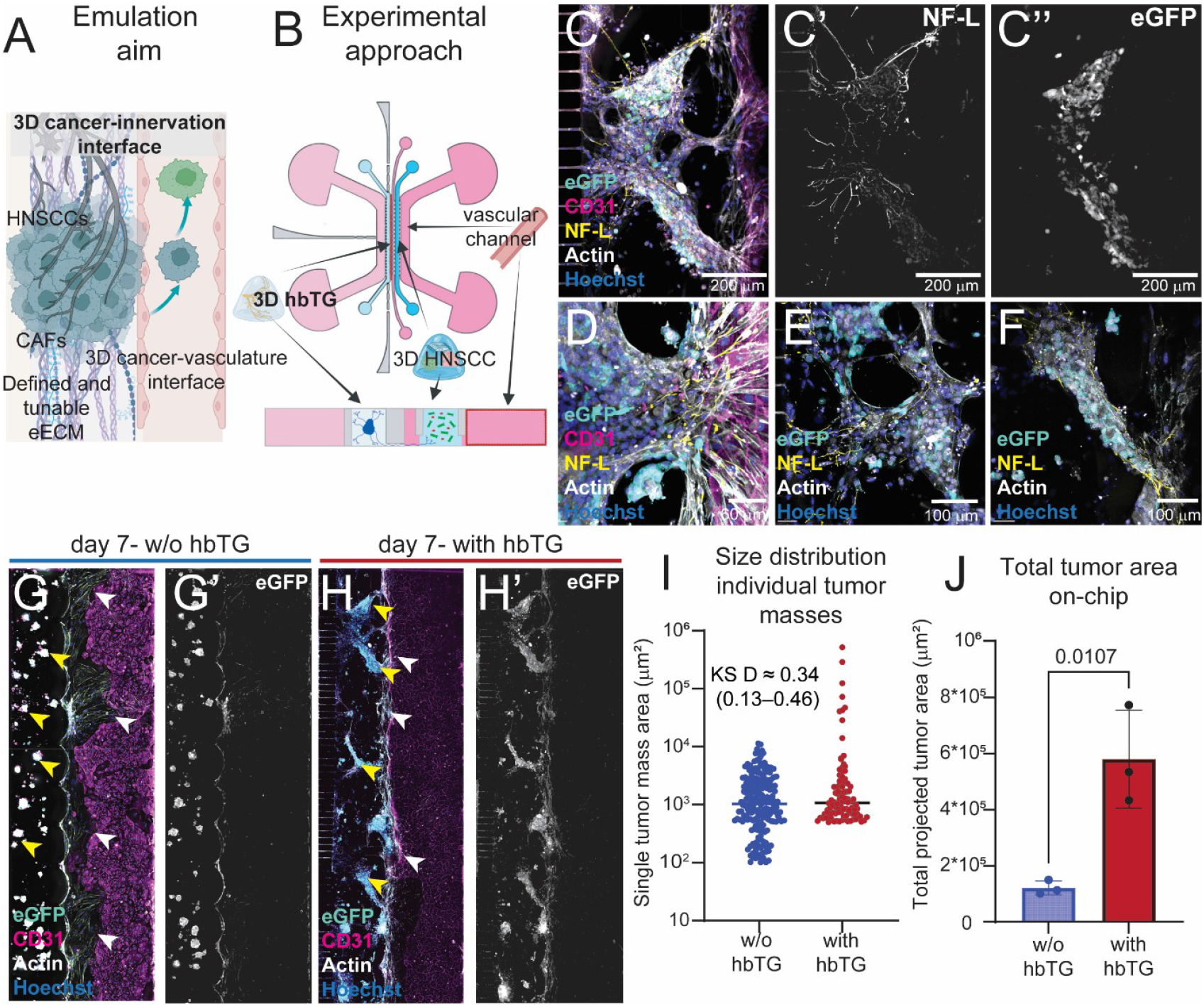
(next page). Complete HNSCC-on-a-chip: 3D trigeminal innervation-3D HNSCC-vasculature interface. **A)** Schematic representation of the emulation aim. **B)** Experimental approach to co-culture3D HNSCC (HCHs: HNSCC cancer cells, CAFs, and HUVECs) at the interface with a HUVEC-lined vascular channel and with hbTG. N = 3 per condition. **C)** Immunofluorescent staining showing HCHs innervated by hbTG at the interface with vasculature. Cyan: GFP/HNSCC; magenta: CD31 (HUVEC); blue: Hoechst3342 (nuclei); yellow: neurofilament light (NF-L); white: actin. **C’**: neurofilament (NF-L) channel of C. **C’’**: eGFP channel of C. **D)** Higher magnification of C, showing in higher detail the innervation of a vascular invasion front. Cyan: GFP/HNSCC; magenta: CD31 (HUVEC); blue: Hoechst3342 (nuclei); yellow: neurofilament light (NF-L); white: actin. **E, F)** Immunofluorescent staining showing innervation of HCHs by hbTG on-a-chip. Cyan: GFP/HNSCC; blue: Hoechst3342 (nuclei); yellow: neurofilament light (NF-L); white: actin. **G-H’)** Immunofluorescent staining showing differential tumor growth in presence (H, H’) or absence (G, G’) of trigeminal innervation after 7 days of culture on-a-chip. Cyan: GFP/HNSCC; magenta: CD31 (HUVEC); blue: Hoechst3342 (nuclei); (NF-L); white: actin. Yellow arrowheads: tumor masses. White arrowheads: vascular invasion fronts. **I)** Size distribution of individual tumor masses when HCHs are cultured with or without hbTG. Box highlights larger tumor masses, observed only in the presence of hbTG-innervation. Chip-level Kolmogorov–Smirnov analysis, KS D = KS distance. **J)** Total projected tumor area per chip. T-test, exact p value is indicated on the graph.

Within 7 days, we observed a radical change in the self-organization of HCHs cultured in the presence of hbTG. While non-innervated HCHs formed small, round shaped spheroid-like clusters (Figure 6G, G’; see also Figure 3), hbTG presence induced the aggregation of HCHs into large tumor structures, often spanning the entire breadth of the 3D HNSCC compartment (Figure 6H, H’). Innervated HCHs formed significantly larger structures (Figure 6I; KS D ≈ 0.34 [range: 0.13-0.46]) and a much larger overall tumor volume (Figure 6J) within the HNSCC compartment than non-innervated HCHs. At the same time however, innervated HCHs tended to invade the vascular channel less than non-innervated HCHs (Figure 6E-F’, white arrowheads).

## DISCUSSION

A central challenge in cancer biology is achieving a holistic and mechanistic understanding of the tumor microenvironment (TME), in which cancer cells interact with extracellular matrix (ECM), vasculature, nerves, and stromal populations under controlled physical and biochemical constraints. Traditional reductionist models have yielded invaluable insights into individual components of the tumor microenvironment, yet they fall short in capturing the behaviors that arise from their integration. In this context, organ-on-a-chip technologies offer human-specific platforms capable of bridging the gap between the fine mechanistic dissection enabled by *in vitro* models and *in vivo* complexity (*3*). The recent decision by regulatory agencies, including the U.S. Food and Drug Administration, to allow, under defined circumstances, clinical testing pipelines that do not rely on animal experimentation further underscores the urgency and relevance of developing predictive, human-based preclinical models (*49, 50*).

In designing the present platform, we focused on three fundamental and broadly conserved interfaces that shape solid tumor behavior: tumor-extracellular matrix, tumor-vasculature, and tumor-innervation. Each of these elements is increasingly recognized as a driver of tumor progression, therapeutic response, and patient well-being (*19, 51, 52*). We thus devised a simple yet powerful design that allows simultaneous emulation and imaging of 3D neural, tumor, and vasculature compartments, in three adjacent channels. The central 3D HNSCC channel enables the study of the behavior of different cells composing HNSCC in an ECM of choice. The direct contact with the adjacent vascular channel enables the study of metastatization via vasculature, as well as the emulation of systemic delivery of drugs, therapeutic immune cells, and other therapeutic approaches. The asymmetric contact with vasculature could enable the study of hypoxia gradients formation (*53*). The direct contact between trigeminal axon terminals and the 3D HNSCC allows fine investigation of the effect of trigeminal innervation on HNSCC behavior. In a complete HNSCC-on-a-chip setup, the presence of a separated trigeminal chamber, connected to the 3D HNCC chamber via microgrooves, allows imaging-based analysis of neuronal activity in response to tumor progression and therapeutic treatments, opening the possibility of studying HNSCC-associated pain-on-a-chip. While immune components are undeniably central to HNSCC and overall tumor biology, their integration into complex microfluidic systems poses additional challenges (*54*) and represents a logical next step, yet beyond the scope of the current work. Similarly, integration of a HNSCC-bone interface, which constitutes a clinically relevant HNSCC invasion route and was recently proposed in an innovative bone-on-a-chip device (*22*), will require further engineering of the platform to properly include a bone compartment.

The extracellular matrix plays a particularly critical role in HNSCC, where extensive clinical and experimental evidence links aberrant ECM composition and mechanics to invasion, treatment resistance, and poor prognosis (*55*). Most existing 3D HNSCC models rely on undefined matrices such as Matrigel® or Myogel, which suffer from batch-to-batch variability and limited tunability, or single-component ECMs, such as collagen and fibrin, which provide limited biochemical complexity (*28, 56*). These limitations hinder mechanistic interpretation and compromise translational relevance. By contrast, fully defined engineered ECMs enable precise control over stiffness, ligand composition, and degradability, parameters that are essential for dissecting how physical and biochemical cues regulate tumor–stroma interactions. Our results support the notion that ECM composition profoundly influences HNSCC organization, invasion modes, and drug sensitivity, and further show that ECM design cannot be a passive choice but a central experimental variable. Specifically, we show that a hyaluronic acid–PEG–collagen– laminin matrix (HPVCL) promotes the formation of dense, discrete tumor masses composed of HNSCC cells, cancer-associated fibroblasts and endothelial cells (collectively: HCHs), closely resembling the compact architecture observed in patient tumors. In contrast, fibrin-based matrices support a more diffuse and dispersed cellular organization. Notably, HCHs embedded in the HPVCL matrix exhibited increased sensitivity to cisplatin compared with those cultured in fibrin, suggesting that ECM-mediated changes in cell state, proliferation, or stress responses may critically modulate chemotherapeutic efficacy. Based on mesh size alone, fibrin ECMs should allow easier diffusion of cisplatin than HPVCL, as the pore size is in the range of micrometers for fibrin ECMs (*57*) and of nanometers for HPVCL eECMs (*58*). This finding highlights the importance of systematically interrogating ECM-dependent drug responses and motivates future studies aimed at linking matrix properties to specific resistance mechanisms (*55, 59–61*).

Vascularization is a second essential dimension of tumor pathophysiology, providing trophic support, waste removal, and metastatic dissemination. By culturing 3D HCHs constructs adjacent to an endothelial cells-lined vascular channel, we observed rapid, robust, directional migration of tumor cells toward the vascular compartment. Whether this behavior is driven primarily by gradients of trophic support or endothelial cells-derived signaling cues remains to be determined. Interestingly, on-chip we did not observe the capillary-like vascular networks within the 3D tumor masses observed in fibrin or HPVCL isolated 3D cultures. As HUVECs survived and formed small vessels in all eECMs tested, it is improbable that they underwent apoptosis when the same eECMs and culture conditions were used on-chip. We rather hypothesize that the HUVECs resuspended in the eECM merged with the main vascular wall lining the vascular channel, during the formation of the endothelial barrier. This is consistent with *in vivo* observations showing that endothelial cells can integrate into the endothelium of injured blood vessels (*62*). When cultured adjacent to an endothelium-line vascular channel, 3D HCHs showed radically different self-organization and migratory patterns depending on the ECM chosen. 3D HCHs cultured in HPVCL self-organized in discrete tumor masses, as observed when they were cultured in simple 3D cultures, and then migrated collectively towards the vascular channel. On the other hand, 3D HCHs cultured in fibrin migrated on several independent paths from the ECM to the vasculature. When cultured for 7 days, HCHs embedded in HPVCL matrix displayed a pronounced dimorphism, with distinct cellular subpopulations emerging near and distant from the vasculature, a phenomenon absent in fibrin-based matrices. This observation reinforces the possibility that ECM composition may regulate the balance and spatial distribution between quiescent, invasive, and proliferative tumor cell states, a hypothesis with important implications for metastasis and therapy resistance (*63*). The response to cisplatin in vascularized chips was attenuated compared with isolated 3D tumor cultures, a finding consistent with emerging literature in vascularized tumor models. While this effect cannot be readily attributed to impaired drug diffusion, as substantially larger molecules readily diffuse from the vascular channel through the 3D ECMs, the delivery purely from the vascular channel potentially increases the barrier effect exerted by the ECM. Furthermore, endothelial cells themselves, via endothelial cells-mediated drug sequestration, metabolic modulation, or pro-survival signaling, may further contribute to the observed protection (*64, 65*). Addressing this question will require future enrichment of the model, for example by incorporating more physiologically complete vessel walls, including pericytes, smooth muscle cells, and potentially tumor-derived endothelial-like cells.

A distinctive feature of the present platform is the incorporation of functional sensory innervation. Growing evidence shows that neural activity is an active regulator of tumor growth and invasion, particularly in HNSCC (*17, 66*), where trigeminal innervation also underlies cancer-associated pain, a major clinical burden and a frequent dose-limiting side effect of cisplatin (*67*). Here, we used human embryonic stem cell to generate 3D bioengineered trigeminal ganglia (hbTG) that form functional, action potential–competent neuronal networks and robustly innervate 3D HCHs. In line with the original protocol, which successfully generated a diverse population of trigeminal neurons and glial cells in 2D, our 3D hbTG contained both neuronal and glial cells and expressed markers typical of diverse neuronal subpopulations, including action potential-conducting nociceptors responsive to ATP. We decided to proceed via pre-differentiation in 2D, followed by encapsulation and maturation in an injectable and fully defined 3D eECM, rather than using well-established sensory organoids (*68*), to minimize damage to neuronal structures when injecting the 3D hbTG in the chip. An unexpected advantage consisted in the abundant sprouting and bundling of axons from the self-organized hbTG through the eECM towards the edges of the 3D construct, reminiscent of *in vivo* patterns of axon sprouting and nerve bundling in the trigeminal complex (*69*). Other designs have been proposed that allow insertion of ganglia and larger organoids within organ-on-a-chip devices without injection (*70, 71*). Yet, we believe that our “fully injectable ganglia” strategy provides a valuable, complementary approach that can significantly expand the possibilities to integrate complex innervation in a variety of new and already existing organ-on-a-chip designs. The use of fully defined eECMs furthermore opens the possibility to systematically assess whether and how eECM properties affect the establishment neuronal and glial cell diversity. Notably, we observed that trigeminal innervation leads to increased growth of the tumor mass and altered invasion dynamics. The mechanisms underlying these effects may involve release of signaling molecules from axon terminals, or bidirectional electrical and biochemical signaling, as well as axons serving as physical scaffolds for tumor growth and potentially metastatization, paralleling observations from *in vivo* studies that support complex effects of innervation on HNSCC progression and metastatization (*66, 72, 73*). Beyond tumor biology, this innervated platform opens the door to future “pain-on-a-chip” investigations, enabling direct assessment of chemotherapy-induced neurotoxicity and analgesic strategies while adhering to the principles of replacement, reduction, and refinement (3R).

In summary, we developed a fully defined, innervated, and vascularized HNSCC-on-a-chip platform that captures key structural and functional determinants of HNSCC pathophysiology. More broadly, this approach provides a versatile framework for studying solid tumor responses to therapy, drug transport across ECM and vascular barriers, and the neural regulation of cancer growth and pain. While integration of immune components remains challenging, the system is already compatible with investigations of CAR-T and CAR-NK cell behavior in complex 3D environments. We anticipate that continued refinement of this platform will contribute to more predictive preclinical testing and to the rational design of therapies that address not only tumor control, but also patients’ quality of life.

## MATERIALS AND METHODS

### Cell culture

Samples from patients with HNSCC were obtained from an established tumor collection (No 416, The National Board of Health and Welfare in Sweden) at the Department of Otorhinolaryngology, Head and Neck Surgery, at the University Hospital of Linköping, Sweden. The Ethical Committee of Linköping approved the collection (approval no. 03-537), and written consent was obtained from the patients. Tumor cell lines (HNSCCcs) and CAFs were established from pretreatment HNSCC biopsies as described in the original publication (*34*). HNSCCcs and CAFs were cultured in DMEM/F12 (Cat. #11320033, ThermoFisher Scientific) supplemented with 10% Fetal Bovine Serum (FBS; Cat. #A5256701; ThermoFisher Scientific), and 100 U/mL of penicillin/streptomycin (Cat. #11320033, ThemoFisher Scientific). When confluent, cells were passaged by incubation in trypsin/EDTA 0-25% (Cat. #10566024, ThermoFisher Scientific) followed by splitting at the desired concentration, without centrifugation steps. Human umbilical cord vein endothelial cells (HUVECs; Cat. #C-12200, Sigma-Aldrich) were cultured in endothelial growth medium (EGM-2; Cat. #C-22211, Sigma Aldrich) supplemented with EGM-2 supplement mix (Cat. #C-39216, Sigma-Aldrich) and 100 U/mL of penicillin/streptomycin on collagen I (PureCol® Type I Collagen; Cat. #5074-35ML, CELLInk)-coated flasks. HUVECs were detached by incubation in Tryple Express (Cat. #12604021, ThermoFisher Scientific) and plated at the desired concentrations. All experiments were performed using HUVECs between passage IV-VIII.

MESP1mCherry/w-NKX2-5eGFP/w human embryonic stem cells (*74*) were kindly provided by Prof. Claudio Cantù (Linköping University). hESCs were cultured in Essential 8 medium (Cat. #A1517001, ThermoFisher Scientific) on vitronectin-coated (0.1 mg/mL in PBS; Cat. #A14700, ThermoFisher Scientific) plastic tissue flasks. When locally confluent, hESCs were detached via incubation in 0.5 mM EDTA, diluted in Essential 8, and split 1:3 to 1:12.

### hESCs-derived trigeminal ganglia

For hESC expansion, T25 cell culture flasks were pre-coated with vitronectin (0.1 mg/mL in PBS) and maintained at 37 °C in a humidified atmosphere with 5% CO_2_ using Essential 8 (E8) medium supplemented with 50 U/mL penicillin/streptomycin, hereafter referred to as “complete E8”. Differentiation of hESCs into trigeminal ganglion (TG) neurons followed a previously established protocol (*41*) originally developed for generating human iPSC-derived trigeminal neurons in 2D culture. hESCs were seeded in complete E8 at a density of 2.5^*^10^5^ cells/cm^2^ on vitronectin-coated flasks. After 24 hours (day 0), the medium was replaced with E6 (Cat. #A1516401, ThermoFisher Scientific) or TeSR-E6 (Cat. #05990, STEMCELL Technologies) supplemented with SB431542 10 μM (Cat. #1614/10, R&D Systems) and BMP4 5 ng/mL (Cat. # 314-BPE-010, R&D Systems) (E6-SB). From day 2 to day 4, E6-SB was further supplemented with CHIR99021 600 nM (Cat. #72054, STEMCELL Technologies) (E6-SBC). On day 5, medium was replaced with E6 supplemented with SB431542 10 μM. After 14 days of differentiation, the resulting trigeminal precursors were detached using 0.5 mM EDTA and resuspended in a hydrogel containing hyaluronic acid, polyethylene glycol (PEG), and human laminin 521 (Cat. #LN521-05, BioLamina) (see section “Hydrogels composition and fabrication” for details on hydrogels composition). Trigeminal precursors were then cultured within the 3D eECM in Neurobasal (Cat. #21103049, ThermoFisher Scientific) supplemented with 2 % B27 (Cat. #17504044, ThermoFisher Scientific), NGF 50 ng/mL (Cat. #450-01-100UG, ThermoFisher Scientific), BDNF 50 ng/mL (Cat. #AF-450-02-10UG, ThermoFisher Scientific; Cat. #78005, STEMCELL Technologies), GDNF 50 ng/mL (Cat. #450-10-10UG, ThermoFisher Scientific; Cat. #78058, STEMCELL Technologies), DAPT 10 μM (Cat. #J65864.MA, ThermoFisher Scientific), and non-essential aminoacids (Cat. #11140050, ThermoFisher Scientific), for up to 40 days to generate human bioengineered trigeminal ganglia (hbTG). For calcium imaging, hbTG were infected with AAV9-CAG-jGCaMP8f-WPRE (2.6^*^E^11^ GC/hbTG; stock titer: 2.6^*^E^13^ GC/ml, Addgene) on day 34 of differentiation in eECM. At day 40, hbTG were treated with ATP 1 mM, and calcium influx was live imaged with a Zeiss CLSM. Videos were analyzed with Fiji/ImageJ (*75*).

### Hydrogels composition and fabrication

#### Hyaluronic acid / PEG-based hydrogels

The following reagents were used to produce Hyaluronic acid (HA) / PEG-based hydrogels: sodium hyaluronate (100-150 kDa HA, Cat. #HA100K-5, LifeCore Biomedical, Minneapolis, USA); 8-arm PEG azide (PEG-Az8, 10 kDa, Cat. #PSB-881-1g, Creative PEGWorks); N-(3-dimethylaminopropyl)-N0 -ethylcarbodiimide (EDC), 1-hydroxybenzotriazole hydrate, N-[(1 R,8S,9s)-Bicyclo[6.1.0]non-4-yn-9-ylmethyloxycarbonyl]-1,8-diamino-3,6-dioxaoctane (BCN-NH2, Cat. #745073-100MG; Merck), 2-(N-morpholino) ethanesulfonic acid (MES) (100 mM, pH 7), human laminin 521 (Cat. #LN521-05, BioLamina), PureCol (3.2 mg/mL, Cat. #5005-100 ML, Advanced Biomatrix). Cyclic RGD (Arg-Gly-Asp) (cRGD-Az) and the protease-degradable crosslinker VPM-Az2 were synthesized as previously described (*76*).

HA was functionalized with BCN (HA-BCN) as follows. HA was dissolved in MES buffer (100 mM, pH 7, 40 mL) on an orbital shaker for 2 hours. 100 mg BCN-NH2 were dissolved in a mixture of acetonitrile and milliQ water (MQ) (5:1 v/v, 6 mL), 1-Hydroxybenzotriazole hydrate (HOBt, 83 mg), and 1-ethyl-3-(3-dimethylaminopropyl) carbodiimide hydrochloride (236 mg, EDC). The acetonitrile solution was added in fractions to the hyaluronic acid solution and dialyzed against acetonitrile (10% v/v) for 24 hours, MQ water for 7 days, and adjusted to pH 6.5. The solution was diluted 1:1 with MQ water and freeze dried for 2 days. Two batches of HA-BCN were used for this work. To determine the degree of BCN substitution to HA backbone (%DS), HA-BCN was resuspended in deuterium oxide and analyzed by ^1^H-NMR (%DS batch4 = 12.33%; %DS batch5 = 15.67%. See Supplementary Materials for ^1^H-NMR spectra).

Hydrogels were prepared by mixing HA-BCN and PEG-Az8 at a 4:1 molar ratio. The total polymer stock concentrations used were 1% (w/v%), 1.5%, or 2%, with the volumes adjusted accordingly to maintain the desired ratios. Hydrogels were supplemented with bioactive functionalization: cRGD-Az (1 mM), Human Recombinant Laminin 521 (9 μg/mL), Murine Laminin (Cat. #L2020, Sigma-Aldrich), Collagen I (16-17 μg/mL, Cat. #A1048301, ThermoFisher Scientific, or Cat. #5005-100 ML, Advanced Biomatrix). Partial protease-dependent degradability was ensured by adding the degradable crosslinker VPM-Az_2_ (2 mg/mL). The formulations of all the eECMs used in this work are reported in Table 1. Cells or trigeminal placodes were resuspended in HA-BCN at the desired concentration; the additional hydrogel components were then added to the HA-BCN-cell suspension, with Az-bearing crosslinking agents added at the end. Loaded hydrogels were seeded on well plates or injected into the dedicated channel on the organ-on-a-chip device and allowed to solidify at 37 °C in the incubator for 1 hour before adding culture medium.

**Table 1.** Hydrogels composition.

| Component | Stock conc. | Final conc. | V[μL] for (50 μL) |
| --- | --- | --- | --- |
| **HA-BCN / PEG-Az8 / VPM-Az2/ COL / LAM (HPVCL)** |  |  |  |
| HA-BCN | 20 mg/mL | 14.24 mg/mL | 35.6 |
| PEG-Az8 | 20 mg/mL | 3.56 mg/mL | 8.9 |
| Laminin | 0.1 mg/mL | 0.5 μg/mL | 0.25 |
| Collagen I | 3.2 mg/mL | 16 μg/mL | 0.25 |
| VPM-Az2 | 20 mg/mL | 2 mg/mL | 5 |
| **HA-BCN / PEG-Az8 / cRGD-Az/ LAM (HPRL - for hbTG)** |  |  |  |
| HA-BCN | 10 mg/mL | 7.2 mg/mL | 36.05 |
| PEG-Az8 | 10 mg/mL | 1.8 mg/mL | 9 |
| cRGD-Az | 0.701 mg/mL | 6.3 μg/mL | 0.45 |
| Laminin521 | 0.1 mg/mL | 9 μg/mL | 4.5 |
| **HA-BCN / PEG-Az8 / COL / LAM (HPCL)** |  |  |  |
| HA-BCN | 20 mg/mL | 15.82 mg/mL | 39.55 |
| PEG-Az8 | 20 mg/mL | 3.96 mg/mL | 9.9 |
| COL | 3.2 mg/mL | 17.6 μg/mL | 0.275 |
| LAM | 0.1 mg/mL | 0.55 μg/mL | 0.275 |
| **HA-BCN / PEG-Az8 / LAM (for injection tests)** |  |  |  |
| HA-BCN | 15 mg/mL | 11.2 mg/mL | 37.33 μL |
| PEG-Az8 | 15 mg/mL | 2.8 mg/mL | 9.33 μL |
| Laminin | 1.5 mg/mL | 0.1 mg/mL | 3.33 μL |
| **Fibrinogen / Thrombin (Fibrin)** |  |  |  |
| Fibrinogen | 40 mg/mL | 20 mg/mL | 25 |
| Thrombin | 10 U / mL | 5 U / mL | 25 |

#### Fibrin-based hydrogels

Human fibrinogen (100 mg/mL in 0.9% NaCl; Cat. # F3879-250MG, Sigma-Aldrich) and thrombin (100 U/mL in 40 mM CaCl_2_; Cat. #T6884-100UN, Sigma-Aldrich) stock solutions were diluted to final working concentrations of 40 mg/mL and 10 U/mL, respectively. With fibrinogen and thrombin on-ice, cells were resuspended in thrombin. The two components were mixed at a 1:1 volume ratio immediately before use to initiate fibrin polymerization. Loaded fibrin hydrogels were allowed to solidify for 30 minutes at 37°C in the incubator, before adding culture medium.

### Nanoindentation

#### Sample preparation

For each hydrogel formulation, three replicate 10 µL domes (*n* = 3) were prepared and soaked in PBS for 2 hours before the measurement. For HPVCL eECM (see Table 1), the components were mixed in the following order: HA-BCN was mixed with the peptides (laminin and collagen), the protease-degradable peptide (VPM-Az2) was added, and then PEG-Az8 was added to initiate cross-linking. For the Fibrin ECM, fibrinogen and thrombin were directly mixed for cross-linking. Complete hydrogels were then crosslinked by incubation for 1 hour at 37**°**C.

#### Measurement

The measurements were performed using a displacement-controlled nanoindenter instrument (Pavone: Optics 11 Life, Amsterdam, The Netherlands. Probe specifications: Serial Number = PV240905, Geo Factor in Air = 2.68, Stiffness = 0.43 N/m, Tip radius = 49.5 μm) and the Optics11 Pavone V1.11.2 Software. Prior to sample measurement, cantilever bending was calibrated by indenting a rigid surface and equating cantilever deflection with probe displacement. The probe was aligned with the center of the hydrogel droplet. For each sample, a matrix scan (5x5) in a 100 μm x 100 μm grid scan was performed. The probe was lifted 500 μm between consecutive points to avoid surface interaction during lateral repositioning. In the indentation step the sample was loaded to a maximum force of 0.2 μN using a piezo actuator at a constant speed of 30 μm/s. The force was held constant for 1 second after the peak load was reached, followed by a controlled unloading phase at the same rate. Each measurement was performed once, with n = 3 replicates of the same hydrogel.

#### Data analysis

The Optics11 Data Viewer V2.7.0 Software was used for data analysis. To extract the Youngs modulus from the force-indentation curves, the Hertzian Contact model was applied. Contact point detection was performed using the Pmax = 90% and Pmax = 20% criteria, defined as the points at which the indentation force reached 90% and 20% of the peak load (Pmax), respectively. Only indentation curves with R^2^ ≥ 0.95 were analyzed to ensure robust fitting. For each scan, data points were averaged to yield representative stiffness values for each tissue type and condition.

### Real time quantitative PCR (RT-qPCR)

3D eECMs containing either HCHs or hbTG were disrupted by incubation with TRIzol (Cat. #15-596-018, ThermoFisher Scientific) combined with manual mechanical disaggregation. RNA was then extracted using the Direct-zol RNA purification kit (Cat. #R2052, Zymo Research) according to the manufacturer’s instructions. The resulting RNA was reverse transcribed into cDNA with the High-Capacity RNA-to-cDNA Kit (Applied Biosystems, MA, USA), using a LifeTouch thermal cycler (Bioer Technology, Zhejiang, China). Real-time quantitative PCR (RT–qPCR) for *VIM, CDH1, CDH2, CD44, EGFR*, and the corresponding housekeeping genes *ACTNB* and *GAPDH* was performed on a QuantStudio™ 7 Flex Real-Time PCR System (Applied Biosystems, Waltham, MA, USA) using TaqMan® Gene Expression Assays (*VIM*: Hs00958111_m1; *CDH1*: Hs01023895_m1; *CDH2*: Hs00983056_m1; *CD44*: Hs01075861_m1; *EGFR*: Hs01076090_m1). Reactions were amplified using the standard TaqMan real-time PCR protocol (Thermo Fisher Scientific, Waltham, USA). β-actin (Hs99999903_m1; Applied Biosystems) and GAPDH (Hs02758991_g1; Applied Biosystems) were used as internal reference genes. The thermal cycling conditions were as follows: 95 °C for 10 min, followed by 40 cycles of 95 °C for 15 s and 60 °C for 60 s. RT-qPCR for all other targets (for primers: see Table 2) was performed on a Light-Cycler 480 PCR System (Roche) and targets amplified using PowerUp SYBR Green Master Mix (Cat. #A25777, ThermoFisher Scientific). The thermal cycling conditions were as follows: 95 °C for 10 min, followed by 40 cycles of 95 °C for 15 s, 55 °C for 10s, and 60 °C for 30 s. Data were normalized to the geometric mean of the reference genes. Relative expression levels were calculated using the ΔCt method or the comparative Ct (ΔΔCt) method and are presented as fold changes relative to the housekeeping gene or to the control sample (see figure legends).

**Table 2.**
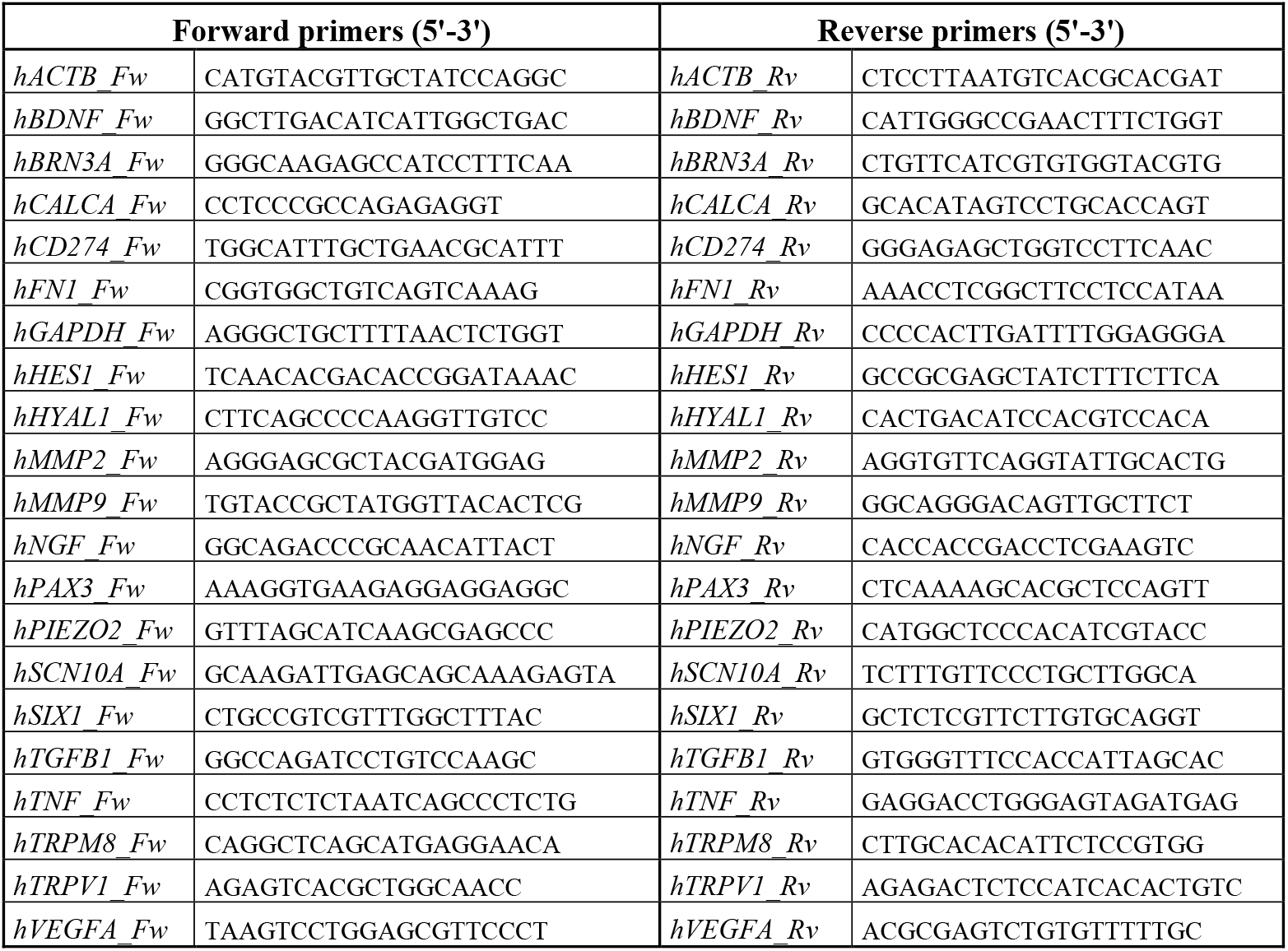
Primer sequences.

### Microfluidic devices design and fabrication

The microfluidic device layout was designed using computer-aided design (CAD) software (AutoCAD®, AutoDesk Inc.). It consists of two main compartments for 3D cell-laden hydrogel cultures, one dedicated to the trigeminal compartment (hbTG compartment) and the other to the target tissue, represented by the 3D HNSCC model (HNSCC compartment). Both compartments are 150 μm high and 700 μm wide, and they are connected by 115 microgrooves (Figure 1C). These microgrooves were specifically designed to allow axonal growth while preventing the migration of neuronal cell bodies. Each microgroove measures 150 μm in length, 5 μm in height, and 10 μm in width, with a 50 μm spacing between them. The hbTG compartment is a single-channel chamber with an inlet (⌀ = 1 mm) for 3D hbTG gel injection and an external channel with two reservoirs (⌀ = 5 mm) for medium supply. Similarly, the HNSCC compartment is a single-channel chamber, featuring gel injection ports (⌀ = 1 mm) to accommodate HNSCCs, CAFs, and HUVECs resuspended in the eECM of choice. The vascular compartment, responsible for supplying medium to the tumoral tissue, includes two reservoirs (⌀ = 5 mm) that serve as inlets for HUVECs seeding. The pillars in the trigeminal compartment have a trapezoidal cross-section, with a maximum length of 200 μm and a minimum inter-pillar distance of 50 μm. In contrast, the target tissue compartment features moon-shaped pillars, measuring 305.1 μm in length with 30.5 μm spacings between them. The pillars’ geometry, size, and spacing were specifically designed to retain the hydrogel upon injection. The moon pillars are specifically intended to achieve the desired level of confined compression (*77*). In order to systematically optimize the two compartments, two models were designed: a complete device and a partial device. The complete design integrates both the target tissue compartment and the innervation compartment.

#### Devices fabrication

The devices are entirely made of PDMS (Sylgard 184; Dow Corning) and consist of two main components: a top microfluidic layer, where the channel structures are engraved, and a flat bottom layer. Their fabrication was carried out using photolithography and soft lithography techniques. In particular, the process began with the creation of a silicon master mold in a cleanroom environment (PoliFAB, Politecnico di Milano) using maskless photolithography. A 5 µm-thick negative photoresist layer (Cat. #SU-8 2005; MicroChem, USA) was spin-coated onto a 4’’ silicon wafer, followed by the polymerization of microgrooves through direct laser writing (Heidelberg MLA100, Heidelberg Instruments). The non-cross-linked photoresist was dissolved using a developing solution (SU-8 developer; MicroChem), the substrate was dried, and two layers of negative photoresist respectively 50 µm and 100µm thick (Cat. #SU-8 2035, Cat. #SU-8 2100; MicroChem, USA) were sequentially spin-coated and cross-linked, resulting in a 150 µm-thick chamber layer. Following the exposure process, the non-cross-linked photoresist was eliminated using the developing solution, leaving behind a negative mold of the desired PDMS microfluidic layer. After curing at 65°C for 2.5 hours, to produce the static chips, the PDMS top layers were peeled off from the resin molds and bonded via air plasma treatment (Harrick Plasma Inc.) onto either a glass coverslip (Ted Pella, 24 × 60 mm) or a flat PDMS layer. Before bonding, holes were created in the top layer using 5 mm biopsy punchers for reservoirs and 1 mm biopsy punchers for the inlets and outlets. The assembled device was then cured at 65°C for 20 minutes to ensure stable bonding. Finally, each device was sealed and autoclaved before use.

#### Technical and mechanical characterization

To ensure that the fabricated PDMS chips met the intended dimensional specifications, key features such as the central channel height, pillars height, and width were analyzed. First, the devices were divided into three sections, with each section containing a single chamber. Each chamber was then further sectioned transversely at the pillar locations to obtain three thin slices. The samples were examined under a microscope, and images were captured using a digital camera. Measurements were performed using Fiji/ImageJ (*75*) and AmScope Camera software (AmScope, UK), with three independent measurements taken per chamber, followed by the calculation of the mean value and the standard deviation.

#### Injectability tests

Once fabricated, the four chip designs underwent an injection test with a functionalized HA-PEG based hydrogel to ensure that the gel could be injected smoothly without lateral leakage. First, 1.5% w/v HA-BCN was mixed with mouse non-functionalized laminin (0.1 mg/ml final concentration) and then with 1.5% w/v PEG-Az8. HA-BCN and PEG-Az8 were mixed at 4:1 ratio. Subsequently the gel was injected into the 3D HNSCC and 3D hbTG channels of the devices and incubated at 37°C and 5% CO_2_. After gelation, a suspension of red polystyrene microbeads (10 µm diameter, Sigma-Aldrich) in phosphate-buffered saline (PBS, Thermo Fisher) was inserted into the medium channels. Brightfield (BF) and phase contrast (PH) images of the channels were captured to assess the presence of the gel-liquid interface. Additionally, videos of beads movement in the medium channels were recorded. Finally, image and video analysis was conducted using Fiji/ImageJ software (*75*).

#### Dextran and rhodamine diffusion tests (Metabolites and Drug Molecules Mass Transport)

On the same microfluidic devices, experimental tests were conducted with molecules of different molecular weights to evaluate molecular transport and barrier permeability. Specifically, the diffusion of Rhodamine B (i.e., 500 Da) and Dextran (i.e., 2000 kDa) across the pillars and microgrooves separating the channels was tested. During the 1-hour experiment, fluorescence microscopy images were acquired at regular intervals and subsequently analyzed using Fiji/ImageJ (*75*).

### Culture of HNSCC-on-a-chip

#### Vascularized HNSCCs on chip preparation

HNSCCs and CAFs were washed with PBS and dissociated using 0.25% trypsin/EDTA for 5 minutes at 37 °C, while HUVECs were treated with TrypLE Express for 4 minutes at 37 °C. The enzymatic reactions were neutralized by adding culture medium (2× the digestion volume), and the cell suspension was then centrifuged (800g, 3 minutes) and cells counted using a Countess 3 Automated Cell Counter (Thermo Fisher). The three cell types (HCHs) were mixed at a 1:3:1 ratio, centrifuged, and resuspended in the hydrogel of choice. For HA-BCN / PEG-based hydrogels, cells were resuspended in HA-BCN to achieve a final cell density of 15 × 10^6^ cells/mL. HA-BCN and cells were then sequentially supplemented with laminin, type I collagen, and VPM-Az2, followed by PEG-Az8. A total of 3 µL of the resulting HPVCL–cell suspension was carefully injected into the 3D HNSCC channel and incubated at 37 °C and 5% CO_2_ until complete gel crosslinking was achieved.

For the fibrin-based condition, HNSCCs, CAFs, and HUVECs (HCHs) were mixed at a 1:3:1 ratio, centrifuged, and resuspended in the thrombin solution at a cell density of 30 × 10^6^ cells/mL. Equal volumes (5 µL each) of the thrombin–cell suspension and fibrinogen solution were combined to generate the fibrin–cell mixture with a final cell density of 15 × 10^6^ cells/mL. 3 µL of this suspension were carefully injected into the 3D HNSCC channel and incubated at 37 °C and 5% CO_2_ until complete gelation occurred.

Upon hydrogel gelation, the vascular channel was first coated with 0.1 mg/mL fibronectin (Sigma Aldrich) dissolved in basal EGM-2 for 1 hour. HUVECs were then seeded into the vascular channel at a density of 15 × 10^6^ cells/ml in complete EGM-2 medium (5 μL of HUVEC cell suspension per vascular channel). Full channel coverage was assessed under a light microscope. The devices were then placed horizontally in a Petri dish (⌀ = 5 cm), in the incubator. From Day 1 on, cells were cultured in a 50% supplemented EGM-2/50% DMEM-F12 mixture, with daily medium changes. For chips loaded with fibrin hydrogels, aminocaproic acid (ACA, 2 mg/mL; Cat. #A7824, Thermo Fisher Scientific) was added to the culture medium from Day 1 to prevent fibrin degradation.

#### Innervated and vascularized HNSCCs-on-a-chip preparation

To generate an innervated and vascularized HNSCC-on-a-chip model, hbTG in HPRL (see Table 1) and HNSCC/CAF/HUVEC in HPVCL (see Table 1) were simultaneously injected into the two adjacent, dedicated channels (See Figure 1). Upon maturation of hESCs into trigeminal placode progenitors (TPPs), the latter were gently detached with 0.5 mM EDTA and resuspended in 1% (w/v) HA-BCN. The HA-BCN / TPP suspension was then supplemented with cRGD-Az_2_ (final concentration: 1 mM), laminin 521 (final concentration: 0.1 mg/mL), and 1% (w/v) PEG-Az_8_ (HA-BCN : PEG-Az_8_ molar ratio = 4:1), and injected as HPRL-hbTG into the “3D hbTG channel” (3 μL/channel). In parallel, HNSCCs, CAFs and HUVECs (HCHs) were resuspended at a total concentration of 20^*^10^6^ cells/mL (ratio = 1:3:1) in 2% (w/v) HA-BCN. The HA-BCN/HCHs suspension was then supplemented with collagen I (final concentration: 16µg/mL), laminin 521 (final concentration: 0.5 µg/mL), and 2% (w/v) PEG-Az_8_ (HA-BCN : PEG-Az_8_ molar ratio = 4:1) 3 µL of this suspension were carefully injected into the “3D HNSCC channel” and incubated at 37 °C and 5% CO_2_ until complete gelation occurred (approximately 1 hour).

Upon gelation, the “vascular channel” was treated as described above to ensure HUVECs adherence, while the “hbTG medium channel” was filled with E6 medium. On Day 1, the medium in the “hbTG medium channel” was replaced with Neurobasal medium supplemented with 2% B27 (Thermo Fisher), MEM nonessential amino acids, 1 mM L-glutamine, 10 μM DAPT, and 50 ng/mL each of nerve growth factor (NGF), brain-derived neurotrophic factor (BDNF), and glial-derived neurotrophic factor (GDNF).

#### Cisplatin treatment on-chip

Following chip seeding, the partial vascularized HNSCC-on-a-chip models were maintained in a 1:1 mixture of EGM-2 and DMEM/F12 without cisplatin on Day 0. On Day 1, the medium was replaced with the same 1:1 mixture supplemented with cisplatin at a final concentration of 3 µg/mL. The cisplatin-containing medium was replenished daily and maintained until Day 4, corresponding to the end of the experiment. Untreated chips, cultured in EGM-2:DMEM/F12 1:1 without cisplatin for the same period of time served as controls.

#### Immunofluorescencestainingonchip

Samples were washed with PBS and fixed via incubation in 4% formaldehyde (Thermo Fisher) for 15 minutes. After three PBS washes, cells were permeabilized with PBS containing Triton X-100 (0.1% in PBS) for 10 minutes at RT. Non-specific binding sites were blocked with Bovine serum Albumin (2% (w/v) BSA; Thermo Fisher) in PBS for 1 hour at RT, after filtering the solution through a 0.2 µm filter to avoid BSA clumps and non-specific fluorescence. Chips were then incubated with primary antibodies (Table 3) overnight at 4ºC. The following day, chips were washed three times with PBS and incubated with the appropriate secondary antibodies (Table 3) overnight at 4°C. After three washes with PBS, chips were incubated for 1.5 hours with CellMask Orange Actin Tracking Stain (1:1000 dilution; Cat. #A57244; Invitrogen, ThermoFisher Scientific) and for 30 minutes with Hoechst 33342 (1:1000; Cat. #1932420, ThermoFisher Scientific). Samples were imaged with a Leica Stellaris 5 Confocal Laser Scanning Microscope and with a Leica DMi8 Widefield Microscope. Finally, images were analysed with Fiji/ImageJ (*75*) and Imaris Viewer (Oxford Instruments).

**Table 3.** List of antibodies used for immunofluorescent staining.

| Target | Cells | Host | Manufacturer | Catalogue # | Dilution |
| --- | --- | --- | --- | --- | --- |
| CD31 | HUVECs | Mouse | Thermo Scientific | PIMA513188 | 1:10-20 |
| Ki67 | Proliferating cells | Rabbit | Thermo Scientific | MA5-14520 | 1:200 |
| Cleaved Caspase-3 | Apoptotic cells | Rabbit | Thermo Scientific | 25128-1-AP | 1:200 |
| NEFL | Trigeminal neurons | Rabbit | Thermo Scientific | MA5-14981 | 1:100 |
| NEFL | Trigeminal neurons | Mouse | Thermo Scientific | 10242123 | 1:100 |
| SOX10 | Glial cells | Rabbit | Cell Signaling | 89356 | 1:100 |
| Target | Fluorophore | Manufacturer | Catalogue # | Dilution |  |
| Rabbit | Alexa fluor 700 | Thermo Fisher | A-21038 | 1:200 |  |
| Mouse | Alexa fluor 647 | Thermo Fisher | A-31571 | 1:200 |  |
| Rabbit | Alexa fluor 568 | Thermo Fisher | A-10042 | 1:200 |  |

#### Tumorsizeanalysis

3D stacks acquired with a Leica DMi8 Widefield Microscope were deconvolved with Huygens and used to generate maximum intensity projections in ImageJ/Fiji. Tumor sizes were quantified for each tumor-on-a-chip device using ImageJ/Fiji and analyzed at the chip level to account for tumor clustering within devices. Total tumor mass was compared between conditions (untreated vs cisplatin, Figure 4; w/o hbTG vs with hbTG, Figure 6) using a t-test. Tumor size distributions were analyzed using empirical cumulative distribution functions (ECDFs). ECDFs were computed separately for each chip and then averaged across the chips per condition. For statistical analysis, each chip was treated as an independent experimental unit. Distributional differences between conditions were quantified using pairwise Kolmogorov–Smirnov (KS) distances computed between all chip pairs (untreated vs cisplatin, Figure 4; w/o hbTG vs with hbTG, Figure 6). KS distances (D) are reported as effect sizes summarizing the maximum separation between ECDFs.

## ACKNOWLEDGMENTS

All imaging was performed with equipment hosted by the Core Facility at the Faculty of Medicine and Health Sciences, Linköping University. The authors thank Dr. Vesa Loitto at the Core Facility at the Faculty of Medicine and Health Sciences, Linköping University for support and guidance in the imaging. The silicon wafer micropatterning was performed at PoliFAB, the micro- and nanofabrication facility of the Politecnico di Milano.

## Funding

Linköping University

Politecnico di Milano

Swedish Research Council (VR) (grant number 2025-03512)

Swedish Research Council (VR) (grant number 2023-04675)

Knut and Alice Wallenberg Foundation (KAW 2021.0186)

European Research Council (101044665 PROTECT)

## Author contributions

Conceptualization: AMP, CS, EW, KR, DA, MR, and PP

Methodology: AMP, CS, AC, SN, MR, and PP

Formal analysis: AMP, CS, EW, GBC, DA, and PP

Investigation: AMP, CS, AJ, TA, LKP, GBC

Resources: MS, DA, MR., PP

Data curation: AMP, CS, EW, and PP

Writing – original draft: AMP, CS, and PP

Writing – review and editing: AMP, CS, EW, KR, EHA, MS, MR, DA, and PP

Visualization: AMP, CS, and PP

Supervision: CS, AC, MS, MR, DA, and PP

Project administration: PP

Funding acquisition: DA, MR, PP

## Data availability

All data are either present in the paper and/or the Supplementary Materials, or available upon request to the corresponding author.

