## Supplementary figures and images for "An Innervated and Vascularized HNSCC-on-a-Chip Model Built on Defined and Tunable Engineered Extracellular Matrices"

### Supplementary Text

#### Supplementary Text

##### HABCN Batch 4

% DS = 12,33%

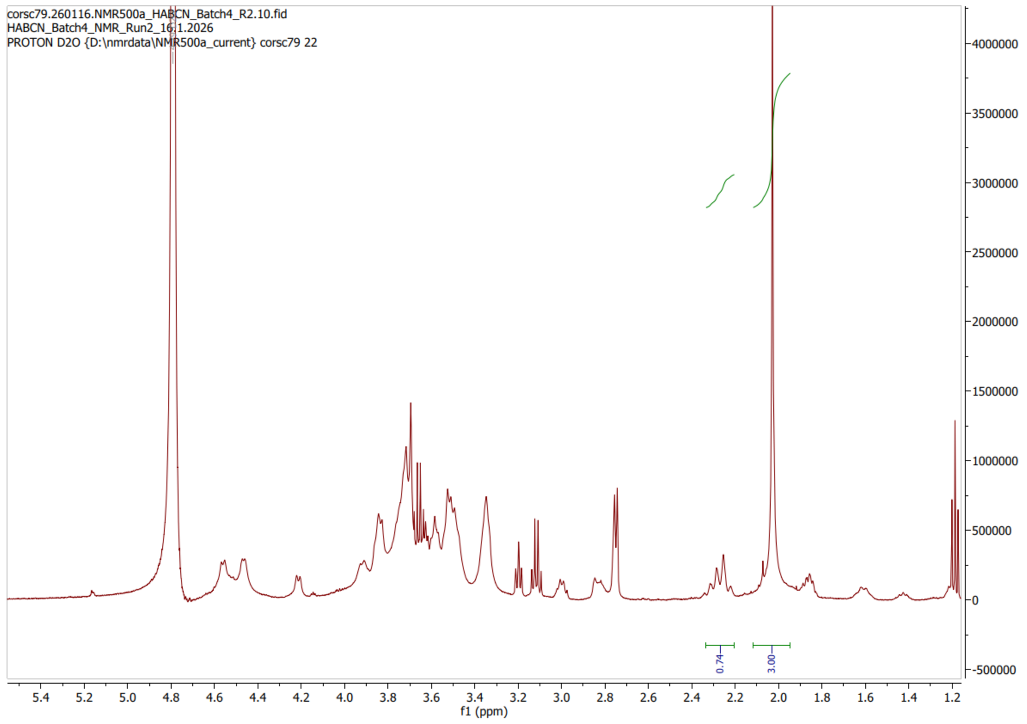


##### HABCN Batch5

% DS = 15,67%

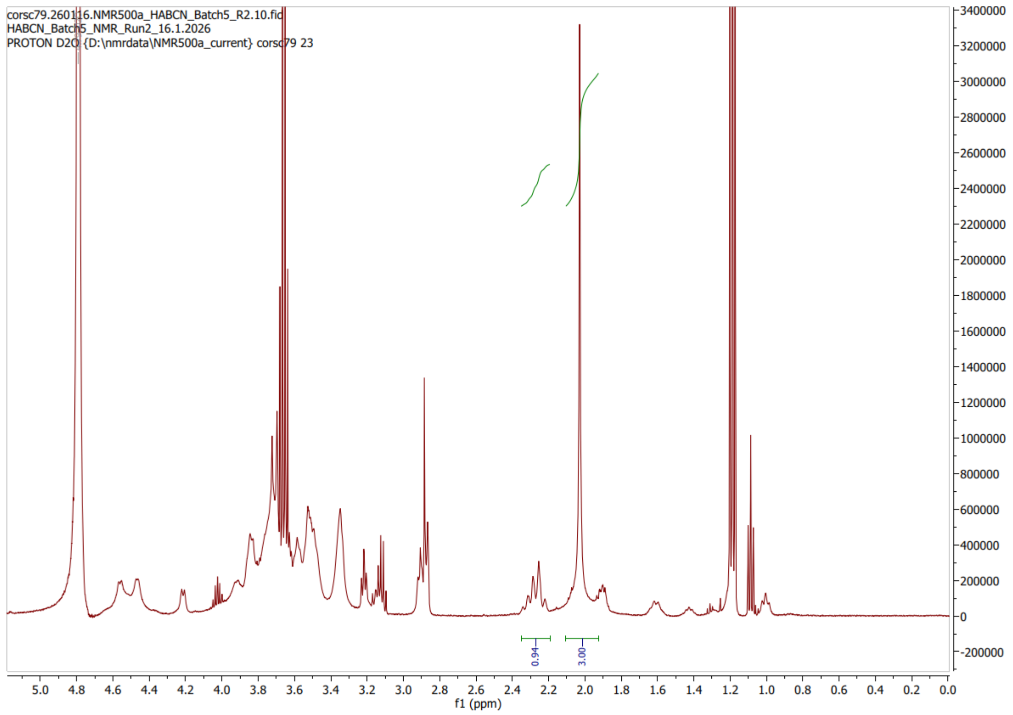
